# Regional Signaling Controls Stem Cell-Mediated Regeneration in an Invertebrate Chordate

**DOI:** 10.1101/2025.09.25.678506

**Authors:** Tal Gordon, Tom Levy, Chester Jiamu Yu, Benyamin Rosental, Lauren Lubeck, Lucia Manni, Irving L. Weissman, Ayelet Voskoboynik

## Abstract

Many tissues harbor quiescent stem cells that activate after injury, yet how local signals regulate this transition is not well understood. The solitary ascidian *Ciona robusta* provides a unique model, as bottom body fragments regenerate while upper fragments fail to do so. By comparing these regenerative and non-regenerative contexts, we reveal striking differences in transcriptional dynamics and signaling environments. Combining flow cytometry, scRNA-seq, transplantation, and fate mapping, we identified a candidate stem cell population with robust proliferative and differentiation potential following transplantation. However, regenerative capacity does not simply reflect stem cell abundance, but instead depends on region-specific signaling cues. Local expression of metabolic, immune and differentiation-related factors further underscores the importance of spatially distinct environments in shaping outcomes. Our findings show how a shared injury response can diverge into regeneration versus failure, highlighting principles that may be leveraged to enhance tissue repair in other systems.

## Introduction

Regeneration, the restoration of tissue following injury or during homeostasis, varies widely across metazoans. While invertebrates like *Hydra* and planarians can regenerate entire bodies, mammals show more limited regenerative ability. In humans, specific tissue types like skin, liver, and blood regenerate well, but others-such as the brain, heart, and spinal cord-do not [1,2]. Variation in regenerative potential may arise from differences in the availability of injury responsive cell types that are able to proliferate and/or differentiate, or from signaling environments that actively restrict or promote regeneration [1,3,4]. Importantly, limited regeneration capacity underlies many diseases, emphasizing the need to understand how different models activate regeneration of diverse tissue types to guide new therapeutic strategies.

Ascidians (Tunicata) are marine invertebrates that represent the closest living relative of the vertebrates and have long been used as a model for developmental and evolutionary studies [5]. One of the interesting features in ascidians is the diversity of regeneration capacity found in closely related species. While colonial species are able to perform whole body regeneration, most solitary species can regenerate only distinct body structures and organs [6]. A comparative approach focusing on ascidian regeneration, examining either species with distinct regenerative capacities or tissues with varied regenerative potential within the same individual, might shed light on the evolution of chordate regenerative capacity and identify key regenerative factors likely conserved in vertebrates.

Circulatory multipotent stem cells were shown to mediate whole body regeneration in colonial species, and a similar mechanism was also suggested to occur in solitary species [6–9]. However, the identity and characteristics of the cell types and signal environment driving regeneration in solitary ascidians remain poorly understood.

Using transgenic animals, we isolated a candidate stem cell population from the solitary ascidian *Ciona robusta* and performed transplantation combined with single-cell RNA sequencing. This approach revealed cells capable of differentiating following injury while maintaining a stem-like subset, enabling characterization of their molecular signature. Leveraging the bidirectional regenerative capacity of this model, we investigated when and how regenerative abilities are established after injury and the involvement of stem cells in the process. Mid-body amputation generates two fragments with distinct regenerative outcomes: a non-regenerative upper fragment and a regenerative bottom fragment [10,11]. In the non-regenerative fragment, the wound remains open and no tissue replacement is observed up to 72 hours post-amputation (hpa). By contrast, the regenerative fragment rapidly closes the wound and restores the missing structures, including the siphons and neural complex, within the same time frame. We hypothesized that by comparing these fragments over time we could reveal distinct cellular and molecular processes that could in turn underlie regenerative ability. Our findings demonstrate that while stem cells are present in both regions, the regenerative outcome is dictated by the local signaling environment rather than stem cell availability. Injury trigger shared early cellular and transcriptomic responses in regenerative and non-regenerative body sections that develop into regeneration specific response at later stages. This study identified and characterized stem cell populations in this model and revealed regeneration-associated factors selectively expressed in regenerative tissues, providing insight into the spatial regulation of tissue repair.

## Results

### Isolation of candidate stem cells in *Ciona robusta*

Isolating stem cells is a key objective in regenerative biology, and achieving this in non-model systems presents a significant challenge [12–15]. In *C. robusta*, a model lacking established stem cell markers, we aimed to enrich for candidate stem cell (cStem) population using ALDH - an enzymatic marker broadly associated with stemness in both mammalian systems [16], [17] and invertebrate species [12,18]. As stem cells are typically small and spherical [12,13], we hypothesized that increased ALDH activity and low granularity might be characteristics of stem cells in *C. robusta.* Using FACS, we sorted three different cell populations based on size and granularity of cells expressing increased ALDH activity (Fig 1A); A population of ALDH positive cells (ALDH), a population of low ALDH activity/high granularity (Control) and a population of a high ALDH activity/low granularity (cStem) (Figs 1A - 1D). To test for enrichment in stem cells markers in our cStem sample in relation to the ALDH and Control samples, we sorted over 10,000 cells from a whole animal, and used the separate populations for single cell RNA sequencing (scRNA-Seq) using 10X Genomics technology. Clustering analysis of the single-cell transcriptomes yielded 20 cell clusters, each expressing a different gene set (Figs 1E-F; S1 and S2 Tables). We annotated each cluster based on gene expression of known markers for the different cell types (Fig 1E; S1 and S2 Figs; S1 Table). Assigning cluster identities based on their FACS-sorted populations and annotated cell-type markers revealed distinct differentiation states, ranging from stem cells in the cStem gate to specialized cells in the Control gate (Figs 1F-G). Cell cycle analysis further differentiated the samples, revealing that the cStem population was enriched for cells in the M and G2 phases, indicative of elevated proliferative activity, whereas the Control population contained a higher proportion of cells in the G1 phase (Fig 1H). Importantly, clusters 0 and 1, were enriched in the cStem sample (Fig 1I) and exhibited strong expression of progenitors and stem-associated genes such as Pou-r (POU6F1) [19], Myc (MYCN) [20], CD34 [21], DDX56 [22–24], ALPL [25,26], and CD53 [27] (Figs 1I and J-K, S1 Table). Although the cStem sample comprises a mixture of clusters, it is notably enriched for progenitors and putative stem cell populations compared to the ALDH and Control samples (Figs 1I-L and M). This enrichment indicates that we successfully identified and isolated an enriched stem cells populations from *C. robusta*, which were subsequently used in the experiments described below.

**Fig 1.**
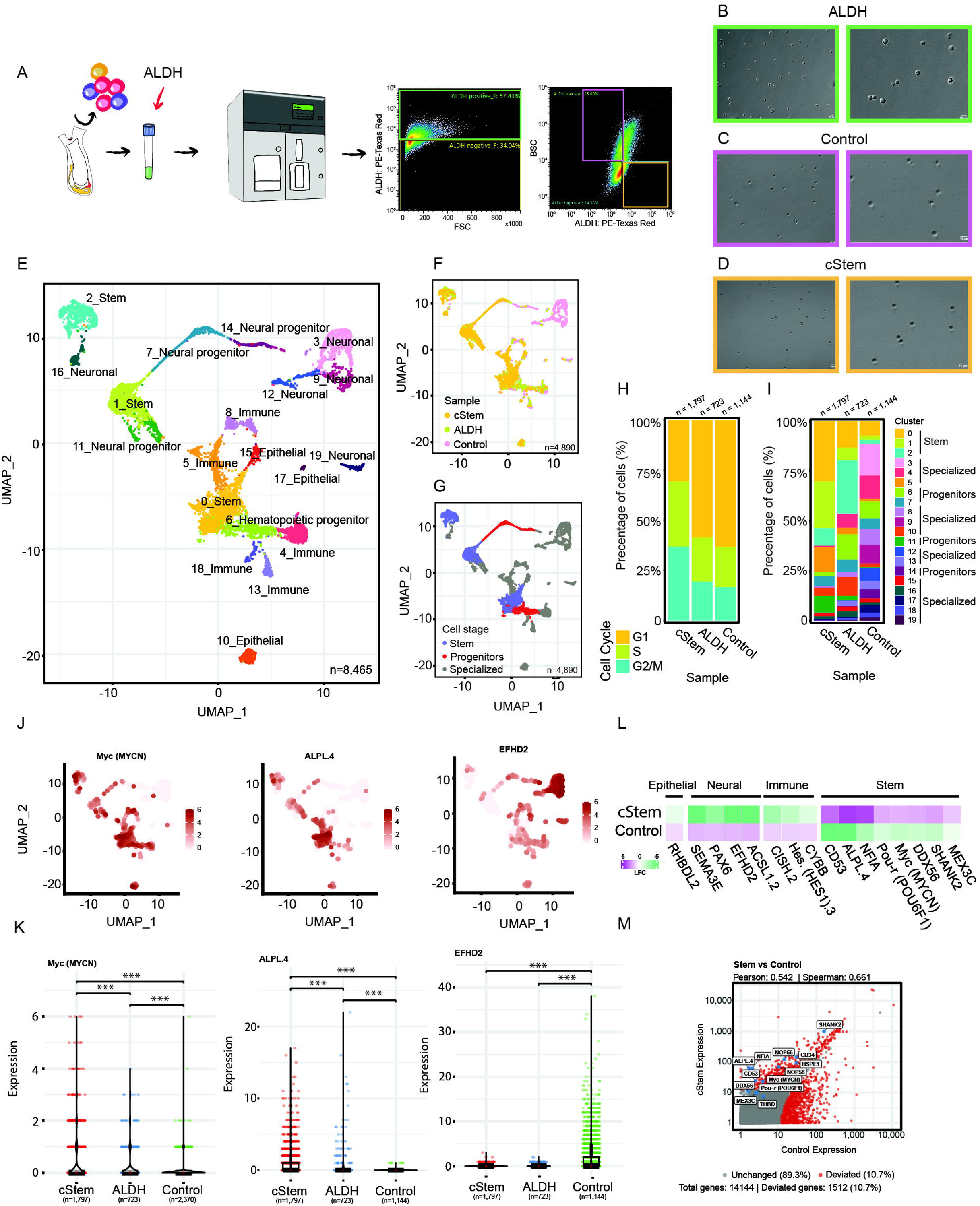
Isolation of enriched stem cells population in *Ciona robusta*. (A) Experimental design for the isolation of enriched stem cells populations in *C. robusta*. Whole dissociated animals were stained with ALDH and sorted into three populations: (B) total ALDH positive cells (ALDH), (C) low ALDH activity/high granularity (Control) and (D) a population of high ALDH activity/low granularity (cStem). (E) UMAP plot with annotated cell type identities based on expressed transcripts of known genes (S1 Table). (F) UMAP plot with annotated clusters based on the FACS gate from which the cells were sorted indicating the clusters differentiation state. (G) UMAP plot showing the clusters differentiation state. (H) Cell cycle analysis of the three populations showing percentage of cells in stage G1, S or G2/M. (I) Percentage of cells originates from different cluster per sample. (J) UMAP expression plots of stem cells markers (Myc and ALPL) and the neuronal marker EFHD2 from scRNA-Seq (S1 Table). (K) Violin plots showing elevated expression of stem cell markers Myc and ALPL in the cStem sample, whereas the neuronal marker EFHD2 is enriched in the Control sample. ^∗^p < 0.05; ^∗∗^p < 0.01; ^∗∗∗^p < 0.001. (L) Matrix plot showing marker gene expression for from scRNA-Seq and the FACS gate they were sorted from (S2 Table). (M) Pseudobulk plot showing stem cell markers expression (in blue) in the cStem sample vs the Control sample.

### Functional validation of candidate stem cells through transplantation and single-cell profiling

A defining characteristic of stem cells is their capacity to differentiate into multiple cell types while maintaining a reservoir of multipotent cells [28]. To investigate the ability of the enriched stem population to engraft and differentiate into diverse cell types, we transplanted cells from donor to recipient of the same parent (to avoid side-effect related to allogeneic rejection [18]) followed by *in vivo* observation and scRNA-Seq of the transplanted cells. Donor animals were transgenic, ubiquitously expressing the GFP reporter from a promoter of the gene Elongation Factor-1 alpha (EF1α::GFP) (see Methods section, Transgenic lines). EF1α::GFP donor animals (n=40) were dissociated and sorted by FACS to generate ALDH-high, BSC-low (cStem) and ALDH-low, BSC-high (control) subpopulations (Fig 2A). Approximately 10,000 cells from each of these subpopulations were then transplanted into WT recipients (Fig 2A). 24 hours post transplantation the animals were injured by amputation of the oral siphon, to encourage a cellular response to the wound. All treated animals survived and regenerated following the injury.

**Fig 2.**
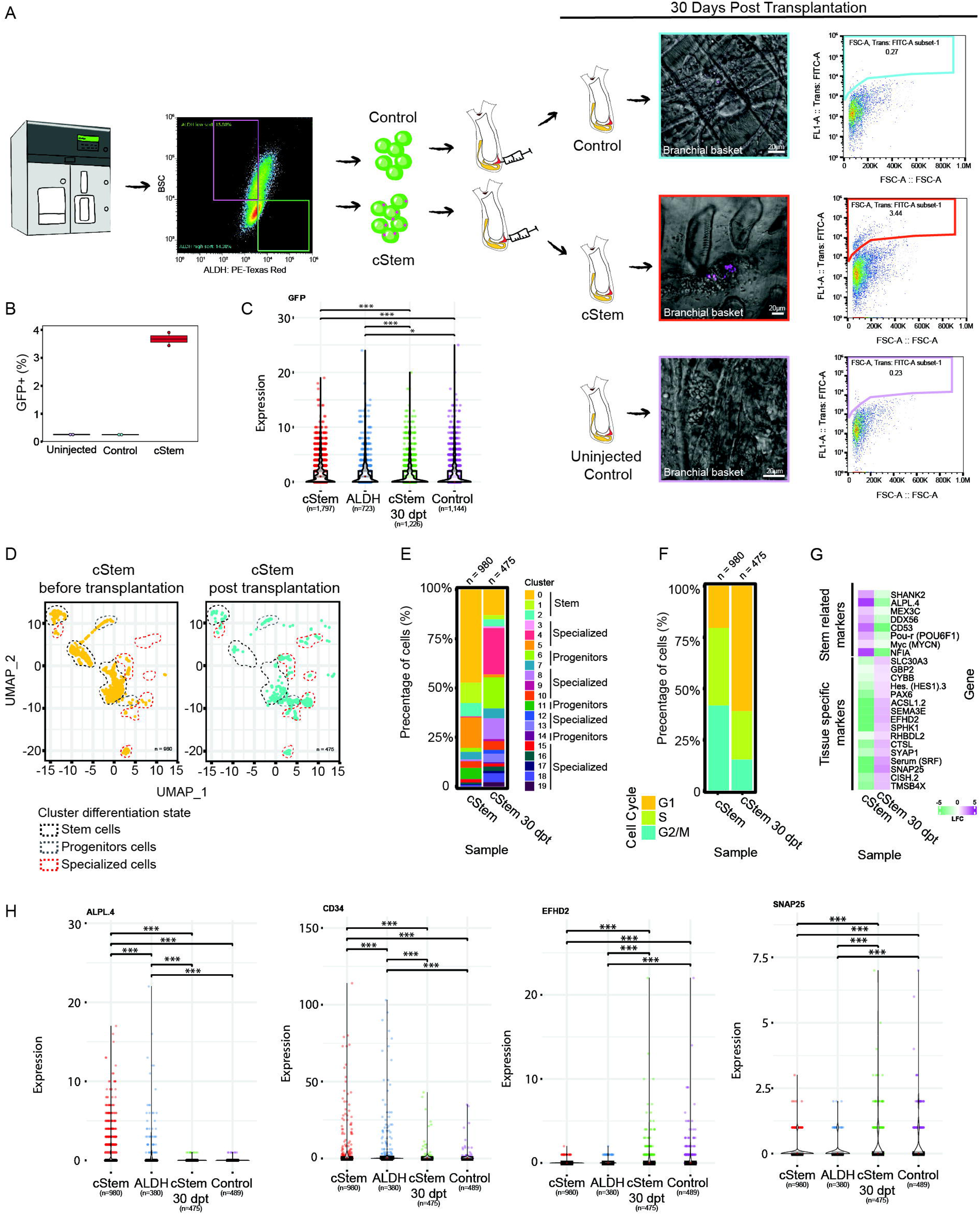
Transplanted enriched stem cell population engrafted and differentiated. (A) Cells from whole GFP+ transgenic animal (n=40) were sorted into two populations: cStem (ALDH+, BSC-) and Control (ALDH-, BSC+). Each population was injected into recipient naïve animals (n=27 Stem, n=15 control). Thirty days post-transplantation, the spatial distribution of GFP⁺ cells within the circulatory system (Branchial basket) of recipient and un-injected control animals was monitored using confocal microscopy and the GFP+ cells were sorted for scRNA-Seq from Stem (n=19) and Control (n=7) animals. (B) The proportion of GFP⁺ cells sorted from recipient animals was higher in the cStem group than in the Control and un-injected groups (n=2 per group of pooled animals). (C) Violin plot of GFP expression levels in the four sample groups. Only GFP expressing cells, representing transplanted cells and excluding host cells, were included in downstream analyses in all four groups to prevent cell type biases. ^∗^p < 0.05; ^∗∗^p < 0.01; ^∗∗∗^p < 0.001. (D) UMAP plot of GFP expressing cells with annotated clusters and the cluster differentiation state showing differentiation of injected cells 30 days post transplantation. (E) Percentage of GFP expressing sorted cells from each cluster in the cStem sample, before and after transplantation. (F) Cell cycle analysis of GFP expressing cells from the cStem sample, before and after transplantation showing percentage of cells in stage G1, S or G2/M. (G) Matrix plot of stem cell and tissue-specific marker expression, organized by the FACS gates from which they were sorted (S1 and S2 Tables). (H) Violin plots showing gene expression in GFP expressing cells 30 days post-transplantation. Stem cell markers: ALPL.4, CD34; differentiated cell markers: CYBB, SNAP25. ^∗∗∗^p < 0.001. (S1 and S2 Tables).

Fluorescence microscopy confirmed a clear GFP signal in the circulatory system of recipient animals thirty days following the transplantation of cStem donor cells. In contrast, recipients of the control cells and un-injected animals showed a low GFP signal (Fig 2A). The relative contributions of donor-derived cStem vs. control cells to recipient animals were assessed by flow cytometry with gating for GFP fluorescent cells thirty days post transplantation. Under identical conditions, cStem donor-derived cells constituted up to 3.44% of recipient cells (n=19 animals, pooled), compared to 0.27% for control donor-derived cells (n=7 animals, pooled). Un-injected controls showed only 0.23% autofluorescent cells (n=7 animals, pooled) (Fig 2A). Averaging the results from two technical replicates, cStem donor-derived cells contributed up to 3.6% of recipient cells (n = 29 animals), compared to only 0.25% for control donor-derived cells (n = 14 animals) (Fig 2B and S3 Table), suggesting the engraftment potential of the injected cStem population. Having established the engraftment potential of cStem cells relative to controls, we performed scRNA-seq to define their molecular signature and differentiation potential. Thirty days after transplantation, GFP^+^ cells were isolated from ALDH high (cStem) recipient animals (n=19), and 3,000 cells were captured for scRNA-Seq. In contrast, control animals (n=7) yielded an insufficient number of GFP+ cells, preventing completion of the single-cell analysis. To ensure that only transplanted GFP⁺ cells, and not autofluorescent host cells, were analyzed 30 days post-transplantation, we assessed GFP expression in the sorted cells and included only GFP expressing cells in all downstream analyses (Fig 2C). Clustering analysis of GFP expressing cells before transplantation (cStem) and 30 days post-transplantation (cStem 30 dpt) suggests a differentiation process, with the cStem 30 dpt sample enriched in differentiated cell populations, whereas the cStem sample was enriched in progenitors and stem cell populations (Figs 2D - 2E). Cell cycle analysis indicated that while the cStem population initially exhibited high proliferation potential, with a large proportion of cells in S/M phases, cStem cells 30 days post-transplantation displayed patterns similar to controls, with a higher fraction of cells in G1 phase, further supporting progression toward differentiation (Fig 2F). Examining the expression of the same stem markers, that were highly expressed in the enriched cStem population prior to transplantation, showed a lower expression of the stem markers (Figs 2G-2H). Flow cytometry of forward scatter (FSC, size) and side scatter (SSC, granularity) 30 days after transplantation showed that cStem donor cells shifted toward a profile resembling the total cell population (S3 Fig). Combined with expression data, this supports differentiation of cStem cells, or their progeny, into multiple cell types within the recipient.

Importantly, the cStem 30 dpt sample retained stem cell clusters, showing that while some GFP expressing cells differentiated, others maintained their stem identity thirty days post-transplantation, supporting that the cStem population is enriched for stem cells. Considering the survival and integration of the enriched cStem population within the recipient 30 days following transplantation, combined with the change in expression pattern of stem markers before and after transplantation indicates that we not only successfully isolated a population enriched with stem cells, but also that those cells were functional enough to differentiate into several different cell types following injury.

### The circulatory system of *Ciona robusta* harbors putative stem cells and progenitors under homeostasis and injury

In *C. robusta*, regeneration has been proposed to be stem cell–mediated, with putative stem cells dispersed throughout the circulatory system without a defined spatial or temporal pattern. Long-distance regeneration is thought to rely on progenitor cells from putative stem cell niches within the circulatory system, particularly the transverse vessels of the branchial basket [9,29] (Fig 3A). Yet regenerative capacity after bisection is spatially restricted, with the upper body fragment failing to regenerate despite retaining parts of the circulatory system and putative stem cells (Figs 3A – 3B). Following isolation and sequencing of the enriched cStem population, we analyzed the expression of 10 marker genes enriched in cStem clusters across regenerated and non-regenerated body sections. Most markers were expressed at comparable levels in both sections, both at 0 hpa and 24 hpa, indicating that stem/progenitor cells are present throughout the body (Fig 3C; S4 and S5 Tables).To analyze the spatial expression of key markers within the two body sections we performed in situ hybridization chain reactions (HCRs) on whole-mount of control (n=3) and regenerating animals (n=3). Our results show ALPL+ cells in the circulatory system at both body parts, in control and regenerated samples (Figs 3D-3E). These observations are consistent with previous studies describing stem cell–mediated regeneration in *C. robusta* and led us to hypothesize that differences in the post-injury signaling environment between body regions drive distinct regenerative outcomes. To investigate this hypothesis, we bisected animals and analyzed proliferation dynamics, immune responses, and gene expression patterns in both regenerated and non-regenerated body regions through the experiments described below.

**Fig 3.**
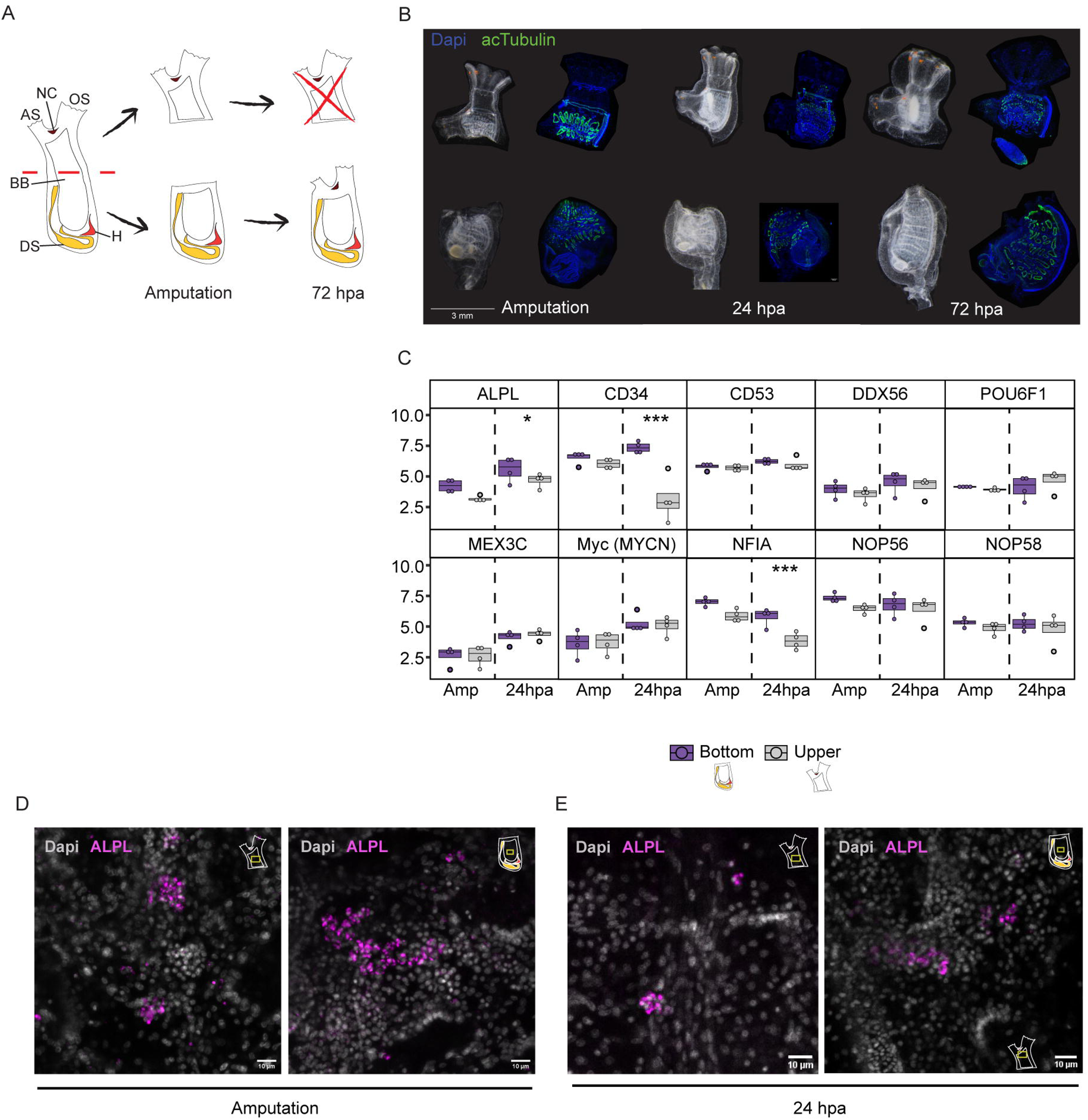
Shared stem cell populations diverge in their response to injury in regenerative and non-regenerative *C. robusta* fragments. (A) Schematic of bisection into two body fragments with distinct regenerative capacities: a regenerative bottom fragment and a non-regenerative upper fragment. Atrial siphon (AS), branchial basket (BB), digestive system (DS), heart (H), neural complex (NC) and oral siphon (OS). (B) In vivo imaging and whole-mount immunofluorescence showing ciliated cells (green, acTubulin) and nuclei (blue, DAPI) in both body fragments following bisection (n = 3 animals per timepoint). (C) Expression of stem cell–associated markers in upper and bottom fragments immediately after amputation and at 24 hours post-amputation (hpa). Box plots show normalized read counts (logCPM) per body section (n = 3 animals per timepoint, n = 4 per region). Data represent the mean. ^∗^p < 0.05; ^∗∗^p < 0.01; ^∗∗∗^p < 0.001. (D–E) Whole-mount immunofluorescence for the stem cell marker ALPL (magenta) reveals widespread localization within the circulatory system of both fragments immediately after amputation (D) and at 24 hpa (E). Insets (top right) indicate the corresponding body fragment and the anatomical region imaged (yellow box).

### Distinct spatial and temporal proliferation patterns following injury in regenerative and non-regenerative tissues

As the onset of cell proliferation often signals the transition from wound healing to regeneration [30], we initially assessed expression patterns of known cell-cycle genes (GLI1 [31], CNBP [32] and KNTC1 [33]) between the different regeneration time points (Fig 4A). Dynamic expression was observed in both tissue types, with consistent upregulation at 12 hpa. These genes were upregulated in the regenerative body part, suggesting a specific response to tissue damage (Fig 4A). To confirm this, we employed EdU labeling to mark cells in the S-phase (Fig 4B and S4A-S4C Figs). Regenerative and non-regenerative body regions were exposed to EdU at different time points post injury (4, 24 and 72 hours) (Fig 4C) and the number of EdU-positive cells was quantified in the regenerating zone (Fig 4D). The regenerative body part showed a higher level of dividing cells in comparison with the non-regenerative body part (Fig 4D). This increase was observed from the earliest time point, 4 hpa, and continued until 72 hpa (Fig 4E). Interestingly, both body parts showed an increase in the number of dividing cells following injury but in a distinct time point. While the regenerative body part showed the highest positive cell count at 24 hpa, the non-regenerative part showed a slower increase which peaked at 72 hpa (Figs 4F-4G). To understand if the increase in dividing cells is local to the injury site or a systemic reaction to injury, the number of EdU positive cells in areas proximal and distal from the injury line was quantified (Figs 4H-4I and S4D-S4J Figs). Both body parts showed a higher number of EdU positive cells in the area proximal to injury line (Figs 4H-4I). At 24 hpa, the regenerative tissue shows an increase in the dividing cells in both proximal and distal areas indicating a systemic response (Fig 4G; S4E-S4F and S4I-S4J Figs). Interestingly, while both body parts showed a constant high level of positive cells at the wound site at all time points, cells that were located farther from the injury site showed a peak in proliferation at 72 hpa and 24 hpa in the upper and bottom body part, respectively (S4F, S4H and S4J Figs). This result might indicate the migration of dividing cells from distal parts of the body to the injured area. This pattern supports our transcriptomics data, reinforcing the notion that injury induces proliferation in both body sections with faster response in the regenerative tissues.

**Fig 4.**
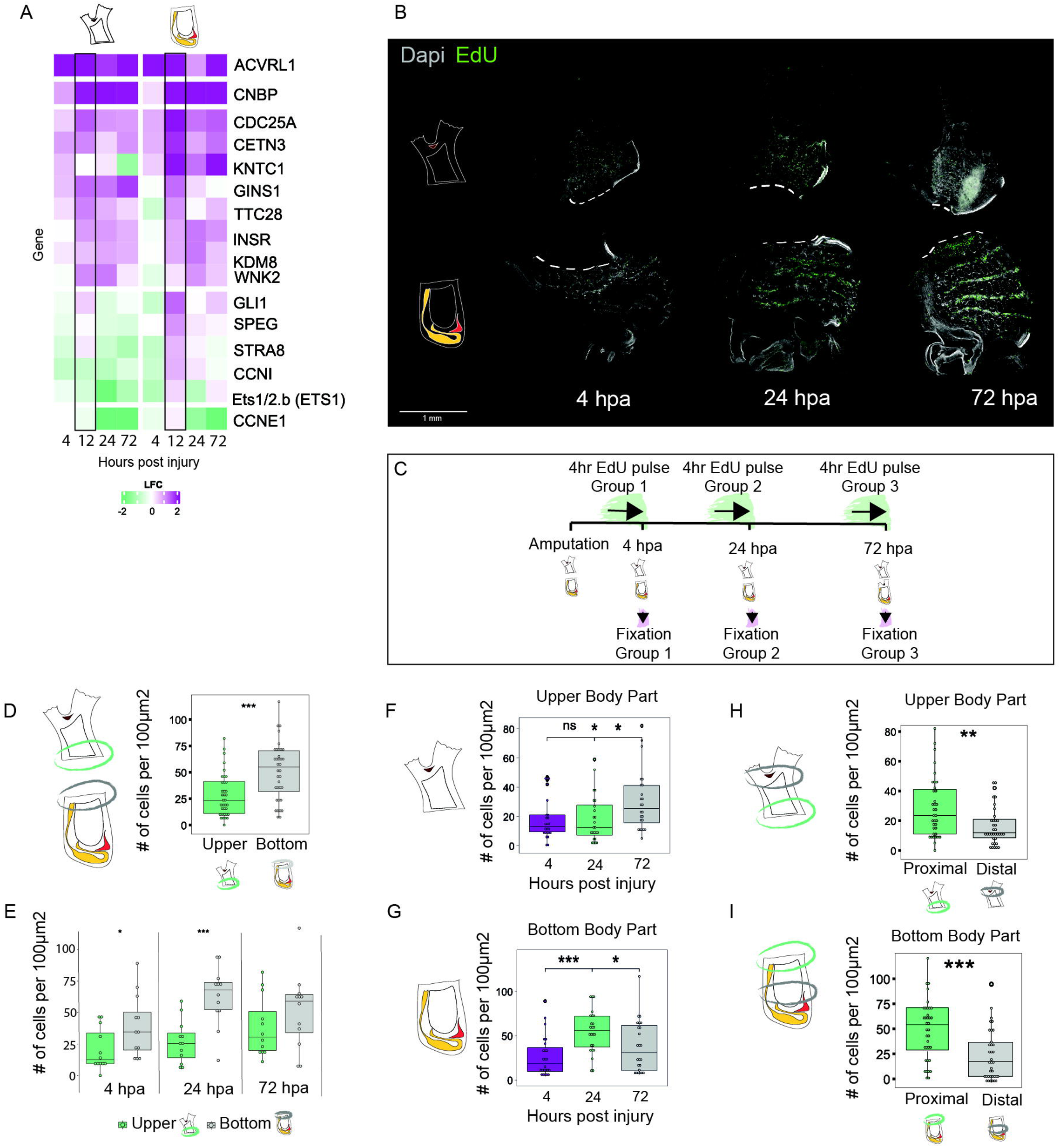
Proliferation dynamics following bisection exhibit distinct temporal expression patterns in regenerative vs non-regenerative tissues. (A) Expression profiling of proliferation-related genes differentially expressed between the non-regenerative upper body part and the regenerative bottom body part. Rows represent genes, and columns show gene expression at each time point. The color scale indicates log fold change. A black frame highlights the 12 hpa time point, where differences between the two body parts are most pronounced. (B) Whole mount immunofluorescence showing proliferating cells (EdU⁺ cells, green) in the two body parts following bisection. White dashed line marks the injury line. (C) EdU experiment set up showing the EdU pulse length and timing. (D-H) Quantification of EdU-positive cells in 100 µm2 sections (n = 3 animals per time point, 4 sections per animal). Box plots display the number of EdU-positive cells in each section. Data are mean. ^∗^p < 0.05; ^∗∗^p < 0.01; ^∗∗∗^p < 0.001. (D-E) Total number of EdU-positive cells near the injury site in both body parts. (D) Combined counts across all time points; (E) counts separated by individual time points post-injury. Two-tailed, t test. (F-G) Number of total EdU-positive cells in the anterior body parts (F) and posterior body part (G) at different time points following injury. Kruskal-Wallis test. (H-I) Total number of EdU-positive cells in tissue proximally and distal from the injury site in the anterior body parts (H) and, posterior body part (I). Kruskal-Wallis test.

### Early Injury-Induced metabolic signaling that supports a regeneration-permissive environment

The widespread distribution of stem cells throughout the animal’s circulatory system, as demonstrated by the HCR of stem cell-related markers, led us to hypothesize that spatial variations in local factors contribute to a regeneration-permissive environment in the regenerative body section. To test this hypothesis, we aimed at identifying differentially expressed genes specifically activated in the regenerative tissue. We conducted a global body-wide differential expression analysis, separately considering each time point (0, 4, 8, 12, 24 and 72 hours post-amputation). To identify stage-specific molecular processes, we then tested for differentially expressed (DE) genes at each time point versus all other time points as well as animal immediately following bisection (0 hpa) (S4, S5 and S7 Tables). Gene Ontology (GO) enrichment analysis identified distinct molecular processes that mark these regeneration stages (Fig 5A; S8-S10 Tables). We observed similar expression patterns of stress-related genes such as HSPA1A[34] and HSPB1[34], along with genes involved in the unfolded protein response (UPR) and ER stress pathways, including CRYAB [35] and USP19 [36] (Figs 5B-5C). Importantly, we found an early regeneration specific expression of metabolic related factors. The bottom regenerative body part showed an increase in GO annotation of pathways related to catabolic process in comparison with the upper, non-regenerative body part (Fig 5D). Furthermore, one of the genes that was upregulated in the regenerative part along all time-points was CASTOR2 [37] a regulator of the TORC1 pathway (Figs 5B and 5E). This pathway regulates cell growth and metabolism by integrating signals from various nutrients and stress cues [38,39]. Additional members of these pathways, such as TSC2 [38,40], PTEN [38,41], PIK3R5 [38] as well as insulin growth factors (IGF1) [42] were also differently expressed in regenerative and non-regenerative tissues following injury (Figs 5B and 5E). GO analysis support a distinct signaling environment between the two body parts, showing enrichment of TORC1 and ERK pathways, key developmental regulators of metabolic processes [39,43], at different time points throughout the regeneration process (Fig 5C).

**Fig 5.**
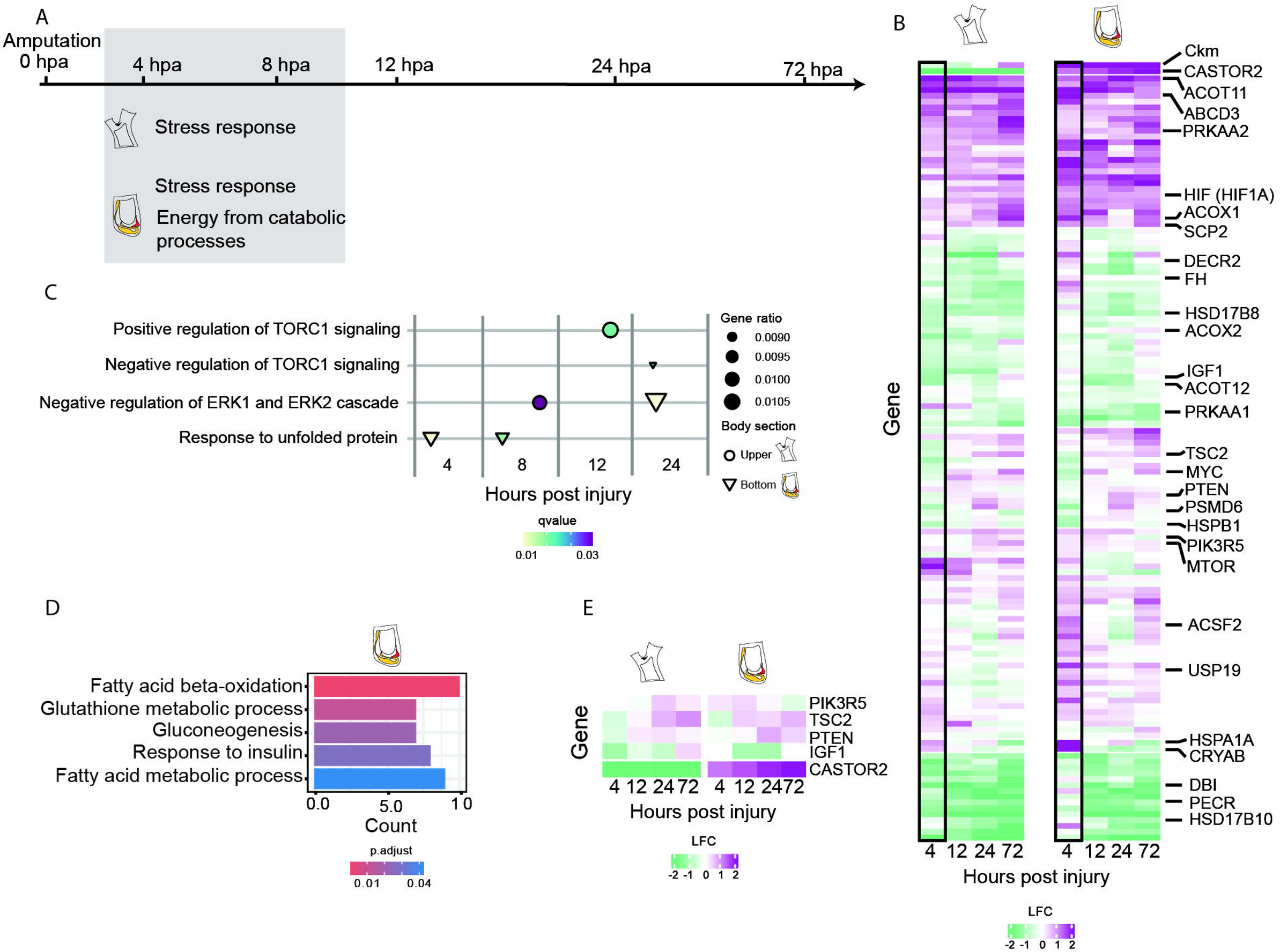
Early regeneration stages exhibit shared stress responses and distinct metabolic activity. (A) Illustration of the experimental timeline showing the main processes occurring at the early time points (4 and 8 hpa) in each body part following an injury. (B) Expression profiling of stress and metabolism related genes differentially expressed between the non-regenerative upper body part to the regenerative bottom body part. Rows represent genes, columns show gene expression in each time point. The color scale indicates log fold change. A black frame highlights the early time point, where differences between the two body parts are most pronounced. (C) Gene Ontology (GO) annotation analysis of both body parts at different time points following injury compared to the immediate post-amputation time point (0 hpa). (D) GO annotation analysis showing metabolic related process upregulating at the regenerative body part (bottom) compared to the non-regenerative body part (upper) at 4 hpa. (E) Matrix plot of log fold change values for TORC1 pathway–related metabolic genes, showing significant differences in expression between the two body parts following injury.

### Late Activation of Developmental Pathways Drives Regeneration Through Extra Cellular Matrix Remodeling and Cell Differentiation

Later stages of regeneration showed a distinct difference between the two body parts (Fig 6A). GO analysis showed that while the upper, non-regenerative, region suffers from starvation and increase in apoptosis, the bottom, regenerative part, showed an increase in glucose metabolism and anabolic processes. These include extra cellular matrix (ECM) remodeling, cytoskeleton organization and cell differentiation (Figs 6A - 6B). Comparing gene expression patterns between the two body parts reveals the different expression pattern of tissue remodeling factors as BMP1 [44,45] and MMP9 [46,47] as well as factors related to growth and differentiation such as SOX6 [48] and HOX genes[49] (Fig 6C). One of the genes that was constantly upregulated in the regenerated body part at all time-points was JAK1 (Fig 6B and S5 Fig; S7 Table). This member of the JAK-STAT pathway plays a crucial role in cytokine-mediated cell proliferation and differentiation [50–52]. To analyze the spatial expression of JAK1 within the two body sections we performed in situ hybridization chain reactions (HCRs) on whole-mount of control, (0 hpa, n=3) and regenerating animals (24 hpa, n=3). Our results show JAK1+ cells in the circulatory system at both body parts, in control and regenerated samples (Figs 6E - 6F). Qualitative assessment suggested a higher number of marker-positive cells in the regenerative bottom body section 24 hours post-injury compared to the non-regenerative upper section, consistent with gene expression data (Figs 6D - 6F). Additional members of this pathway, as well as key members of the MAPK cascade, which act as a central regulator of cell survival, proliferation, and differentiation [53,54], showed a distinct expression pattern between the two body parts (Fig 6D). Together, the expression of these developmentally related factors establishes distinct molecular environments in the two body parts, leading to divergent regenerative outcomes.

**Fig 6.**
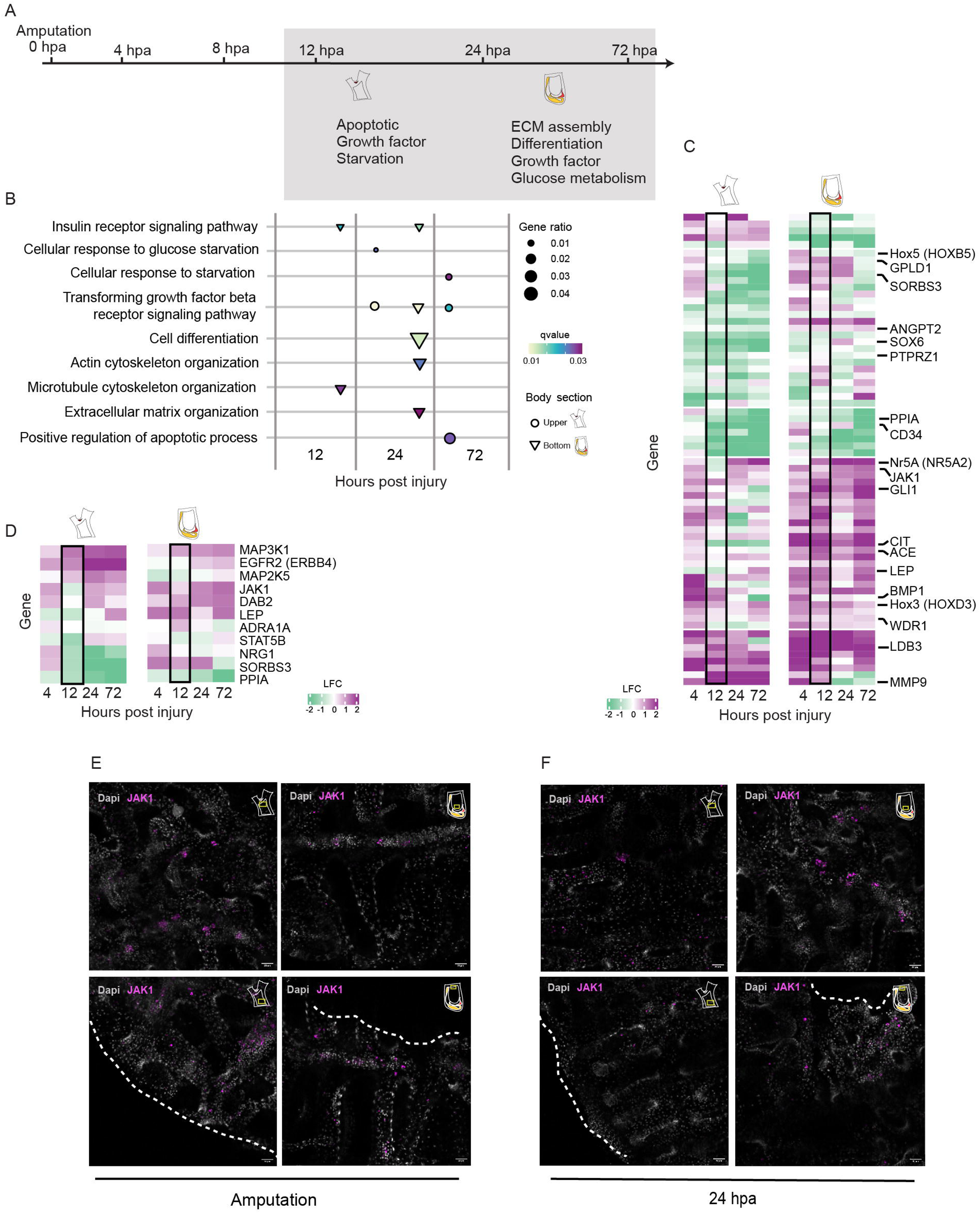
Late regeneration stages exhibit distinct developmental and metabolic activity. (A) Illustration of the experimental timeline showing the main processes occurring at the late time points (12, 24 and 72 hpa) in each body part following an injury. (B) Gene Ontology (GO) annotation analysis of both body parts at different time points following injury compared to the immediate post-amputation time point (0 hpa). (C) Expression profiling of cell differentiation and ECM organization related genes differentially expressed between the non-regenerative upper body part to the regenerative bottom body part. Rows represent genes, columns show gene expression in each time point. The color scale indicates log fold change. A black frame highlights the late time point (12 hpa), where differences between the two body parts are most pronounced. (D) Matrix plot of log fold change values for JAK-STAT and MEPK related genes showing significant differences between both body parts following injury. (E-F) Whole-mount immunofluorescence showing JAK1 expression (magenta) in the circulatory system of both body fragments immediately after amputation (E) and 24 hours post-amputation (F). A higher number of JAK1-positive cells is observed in the posterior body part at 24 hpa compared with the anterior body part (n = 3 animals per timepoint). Insets (top right) illustrate the corresponding body fragment and approximate location of the imaged region.

### Phagocyte cell dispersal follows similar temporal patterns in regenerative and non-regenerative tissues

Tissue regenerative capacity is strongly influenced by the onset of inflammation, a key component of the innate immune response [55]. Inflammatory cells, including phagocytes, perform multiple roles at the wound site, such as clearing damaged tissue and secreting chemokines, metabolites, and growth factors that support regeneration [56,57]. To elucidate the response of the immune system following injury in regenerative and non-regenerative tissues we evaluated phagocytic dynamics, using the PhRodo bioparticles assay [58,59]. These particles became fluorescent following phagocytosis and can thus mark phagocyte cells in vivo. Injection of the bioparticles into the heart of *C. robusta* resulted in wide dispersal of positive cells within the circulatory system (Figs 7A - 7D). Animals were bisected 24 hours post PhRodo injection and the location of the positive cells was monitored at three time-points following injury (4, 24 and 72 hours) and compared between both body parts (Figs 7A, 7C and 7D; S12 Table). Interestingly, no significant difference was found in the number of positive cells between the two body parts, and a similar temporal expression was found with an increase in positive cells at 24 hpa followed by a sharp decrease at 72 hpa (Figs 7E-7G). Phagocytosis related factors such as PAK1[60], PTK2 [61] and IRF8 [62] showed a consist upregulated expression pattern in both body parts following injury (Fig 7H) supporting our in vivo results and suggesting increase in phagocytosis activity following injury regardless of the tissue regenerative potential. This prompted us to examine the expression of immune-related factors to identify signals that may instruct phagocytes to adopt pro-inflammatory and pro-regenerative roles in the different body parts. Comparison of differentially expressed genes between the two body parts at 12 hours post-amputation revealed an upregulation of key regulators of inflammatory responses at the regenerative part, including NLRP3 [63], SMPDL3B [64], NFE2L2 (NRF2) [63,65,66], NR1H4 [67], and BIRC3 [68] (Fig 7I). Together, these findings suggest that although phagocytic cells exhibit similar temporal dispersal patterns in both regenerative and non-regenerative tissues following injury, they may perform distinct roles in each body part, potentially shaping the overall regenerative capacity of each region.

**Fig 7.**
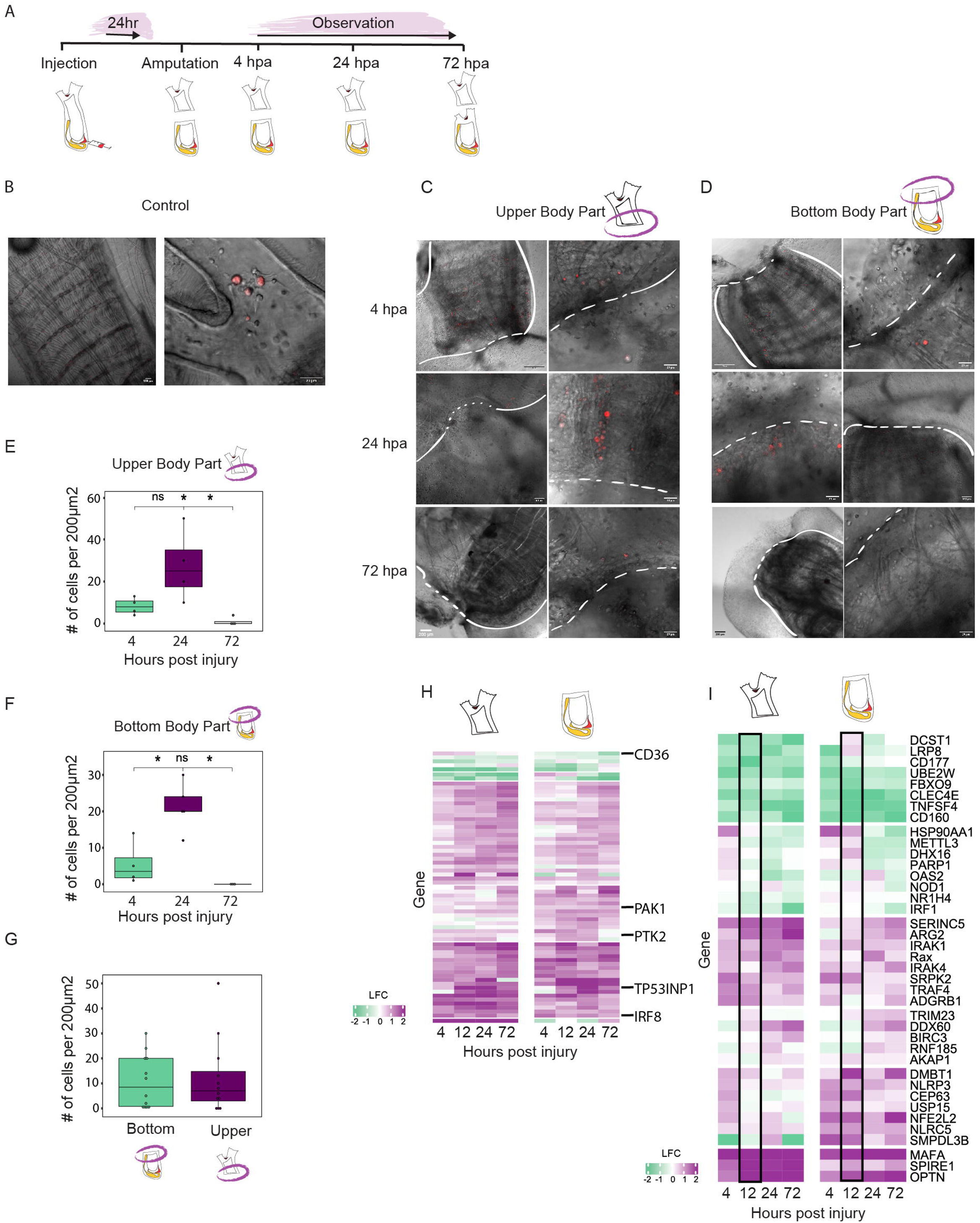
Expression and localization of phagocyte markers following bisection reveal similar temporal patterns in regenerative and non-regenerative tissues. (A) PhRodo phagocytic assay setup and experimental timeline. (B-D) Whole-mount immunofluorescence showing PhRodo⁺ cells (red) in a control uninjured animal, highlighting positive cells in the circulatory system (B), and in the two body parts following bisection (C-D). Right panels in (C-D) show enlarged views. White dashed line represents the injury line. (E-G) Quantification of PhRodo-positive cells in 200 µm2 sections (n = 2 animals per time point, 2 sections per animal). Box plots display the number of PhRodo-positive cells in the area close to the injury site in non-regenerative upper body part (E) and regenerative bottom body parts (F). Data are mean. ^∗^p < 0.05, Kruskal-Wallis test. (G) Number of total PhRodo-positive cells in the area close to the injury site for each body parts. Kruskal-Wallis test. (H-I) Expression profiling of Phagocytosis related genes (H) and innate immune-related genes (I) differentially expressed between the non-regenerative upper body part to the regenerative bottom body part. Rows represent genes, columns show summary of gene expression in each time point. Color scale shows the log fold change. A black frame highlights the late time point (12 hpa), where differences between the two body parts are most pronounced.

### Deconvolution analysis reveals differential stem cell dynamics in regenerative and non-regenerative body fragments

To further resolve the involvement of candidate stem cell populations identified in the cStem dataset during regeneration, we performed reference-based deconvolution of bulk RNA-seq data using our single-cell atlas. This approach enabled inference of cell-type composition over time in both upper and bottom body fragments. Immediately following amputation (0 hpa), both fragments exhibited similar inferred contributions of candidate stem populations, consistent with their broad distribution throughout the animal (Fig 3C and Fig 8). However, following injury, the two fragments diverged markedly in their cellular dynamics. The regenerative bottom fragment displayed pronounced temporal shifts in cell-type composition, characterized by early changes in stem-associated populations, followed by an increased representation of progenitor populations and a subsequent enrichment of specialized cell types over time (Figs 8A - 8D). In contrast, the non-regenerative upper fragment showed relatively limited changes in the inferred proportions of these populations, remaining largely static across time points (Fig. 8A - 8D). Together, these results indicate that while both fragments initially contain comparable stem cell pools, only the regenerative body part supports coordinated transitions in cell states.

**Fig 8.**
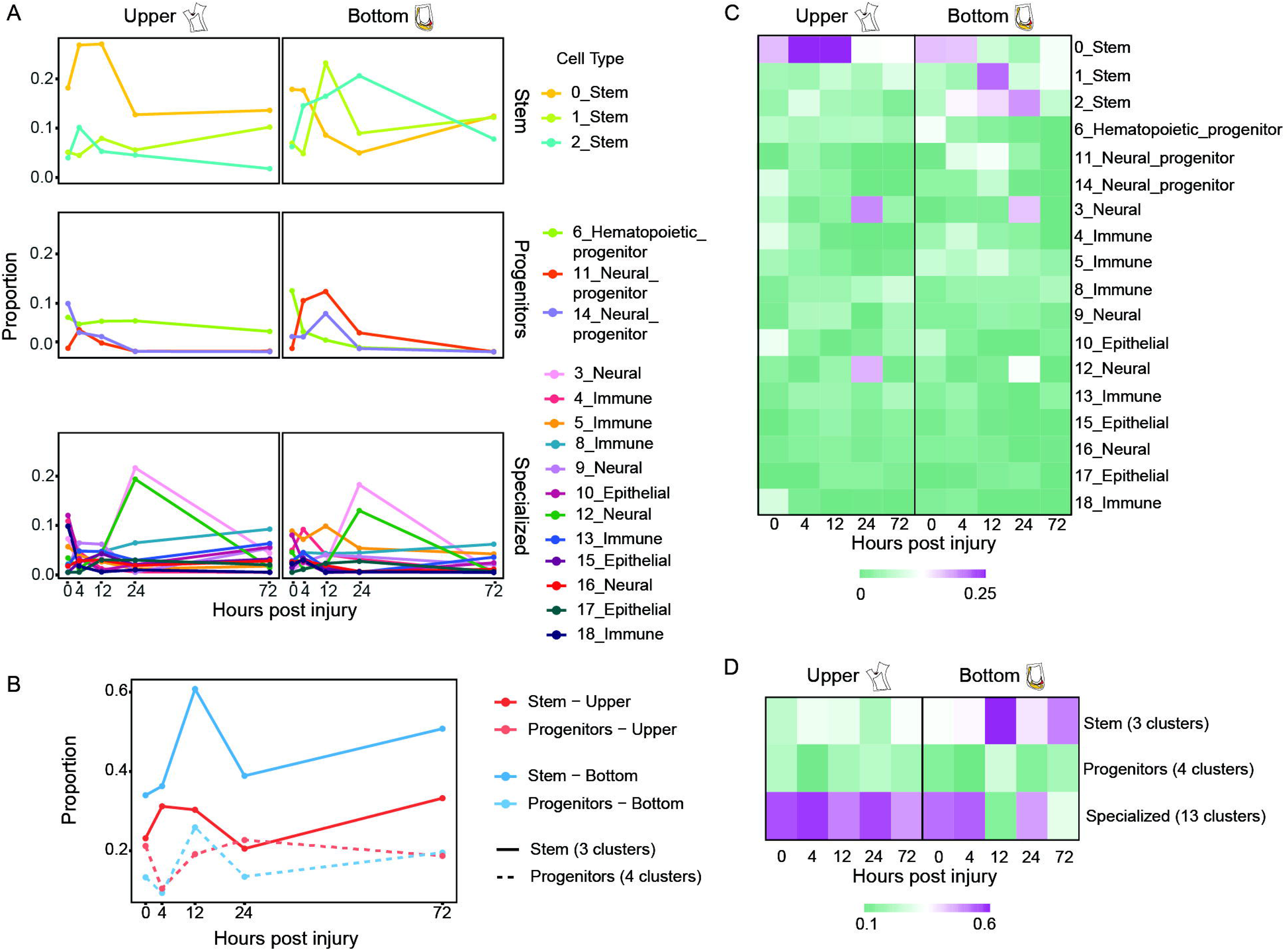
Integration of single-cell and bulk transcriptomics reveals environment-driven stem cell dynamics. (A–D) Deconvolution of bulk RNA-seq data using a single-cell RNA-seq reference to infer cell population dynamics following bisection. (A) Line plots showing temporal changes in inferred proportions of individual cell types in upper and bottom fragments. The y-axis represents the proportion of cells belonging to each of the 17 cell types. (B) Line plot showing aggregated cell states (stem and progenitors), representing the summed contribution of cell types within each category. The y-axis represents the proportion of cells belonging to each of the 2 aggregated states. (C) Heatmaps showing inferred proportions of individual deconvolved cell types across time points. Color represents the proportion of cells belonging to each of the 17 cell types. (D) Heatmaps showing aggregated cell states (stem, progenitor, and specialized), representing the summed contribution of cell types within each category. Across analyses, rows represent inferred cell populations and columns represent time points; color intensity indicates relative contribution to the total cellular composition.

## Discussion

### Establishing method for stem cell isolation in *C. robusta*

Ascidians diverse regenerative potential and close phylogenetic relations to vertebrates offer a powerful system for investigating conserved mechanisms of stem cell-mediated regeneration [7,8,13].

Here we applied a non-species-specific approach to isolate a stem cell-enriched population in *C. robusta*. We validated the presence of putative stem cells in our sample using single-cell RNA sequencing and functional transplantation assays. The isolated cStem population was enriched in clusters expressing stem cell and progenitor markers. Notably, a subset of these cells persisted for 30 days post-transplantation, with some maintaining their stem identity while others transitioned to a tissue-specific state, expressing markers of differentiated cell types, demonstrating both their engraftment and functional integration.

The cStem population represents an enrichment for candidate stem cells rather than a pure stem cell population. It comprises a mixture of stem and progenitor cells, indicating that at least a subset retains the potential to differentiate into other cell types. Single-cell transcriptomic data revealed clear enrichment of stem cell-associated markers in clusters 0, 1, and 2, supporting the presence of stem cells within this heterogeneous population.

Furthermore, GFP expressing cells within the cStem samples retained their stem identity in the recipient body thirty days post-transplantation, further confirming their stem cell quality. The approach developed here for isolating an enriched stem cell population opens the door to a broader range of questions, including the characterization of stem cells across distinct stages of regeneration and comparative analyses among species with varying regenerative capacities. Notably, because the isolated stem cell population was derived from the whole animal, it remains heterogeneous, encompassing multiple lineages and cell types. Applying this method to specific tissues could enable the identification of tissue-resident stem cells, further refining our understanding of lineage-specific regenerative mechanisms.

### Regional and temporal differences in cell proliferation underlies divergent regenerative outcomes

Successful regeneration relies on precise spatial and temporal regulation of mitotic activity following injury [30]. In *C. robusta*, injury induced systemic proliferation in both body regions, with accumulation of proliferating cells at the wounded area (Fig 4 and S4D – S4J Figs). Similar systemic acceleration in proliferation post injury was found in other vertebrate [69] and invertebrate [70] model systems and its function remains unclear [30]. By contrast, the local accumulation of proliferating cells at the wound site, forming a blastema, is a well-established feature across diverse regenerative organisms [71,72]. The cellular sources of blastema cells, however, vary between these systems, ranging from stem cells in planarians [15] to dedifferentiated cells during salamander limb regeneration [73], or a combination of both depending on the injured tissue, as observed in zebrafish [74]. In *Ciona*, the blastema has been proposed to arise from resident stem cell populations [29,75,76]. Our results indicate that although both body regions exhibit increased proliferation at the wound site, consistent with blastema initiation, the upper region fails to close the wound or regenerate, suggesting that these blastemas are non-functional and incapable of restoring lost cell types. Furthermore, the regenerative body section of *C. robusta* showed a higher proliferation rate with overall higher number of dividing cells following injury. While this body part showed the highest positive cell count at 24 hpa, the non-regenerative part showed a slower increase which peaked at 72 hpa. Interestingly, while the level of dividing cells was constantly high at the injury site at all time points, distal cells showed a peak in proliferation at 72 hpa and 24 hpa in the upper and bottom body part, respectively (S4G – S4J Figs). This result might indicate that following injury, a responsive cell divides and migrates to the injured area [30,77]. The differences found in both body sections of *C. robusta* might indicate differences in the temporal control of proliferation, resulting in faster proliferation in the regenerative body part, which in turn could shift the trajectory of tissue repair.

### Distinct signaling environments regulate stem cell-mediated regeneration in a spatial-specific manner

Regenerative capacity is orchestrated by a tightly regulated sequence of molecular events that govern the proliferation and differentiation of specific cell types to restore lost tissues. Our findings indicate that *Ciona robusta* possesses populations of stem and progenitor cells that differentiate following injury and are broadly distributed throughout the body. This observation is consistent with previous injury-based studies in this model, which demonstrated that both anterior organs (e.g., siphons and neural complex) and posterior organs (e.g., the heart) can undergo regeneration, likely through the activation of resident stem cells and progenitor populations [9,29,76,78]. However, following bisection, we observe a striking loss of regenerative ability in the anterior body fragment.

While the bottom fragment regenerates all missing structures within 72 hours, the upper fragment fails to initiate repair and ultimately undergoes degeneration. Our findings indicate that this regenerative failure is unlikely to result from an uneven distribution of candidate stem cells [79,80]. Instead, regional differences in the signaling environment appear to underlie the divergent regenerative outcomes.

An additional factor may be the distinct organs retained by each fragment. The anterior portion lacks the digestive system and heart, whereas the posterior retains them.

Consequently, regeneration failure in the anterior may arise not only from a non-permissive local environment but also from systemic limitations linked to the absence of life-sustaining organs. Our hypothesis is that the signaling milieu differs between anterior and posterior regions, and while structural differences may contribute to these distinct environments, the outcome is the same: a context that does not support regeneration. Future experiments isolating injured tissues from their parent fragments will allow us to directly test whether regeneration potential is governed primarily by local cues or systemic support.

Studies in highly regenerative models that rely on pluripotent stem cells for regeneration, such as planarians and cnidarians, have shown that the signaling environment is critical for successful regeneration, as it regulates stem cell activation and directs differentiation toward the appropriate cell types [81–83]. In the planarian *Dendrocoelum lacteum*, posterior body fragments fail to regenerate a head and ultimately die due to defects arising during early head specification, which depends on the inhibition of canonical Wnt signaling [81].

To investigate whether differences in the signaling environment between body fragments of *C. robusta* dictate regenerative outcomes, we integrated cellular analyses with bulk transcriptomic profiling across multiple time points and anatomically distinct regions to define the molecular programs driving regeneration.

Our findings support a model in which regenerative outcome is specified early following injury. While wounding elicits a conserved response across tissues, it also initiates a region-specific transcriptional program. This spatially distinct response proceeds at later regeneration stages, leading to region-specific regenerative outcomes.

In *C. robusta*, early regeneration stages show a similar stress response in both body regions (Fig 5). Similar results were also found in other vertebrate[84] and invertebrate[85] models, including ascidians [86]. Interestingly, a comparative analysis between the regenerative and non-regenerative body section revealed striking differences in metabolic related pathways (Fig 5). Cellular metabolism is essential for all biological processes and undergoes dynamic changes during regeneration [87,88]. To support proliferation and tissue rebuilding, the cells must rewire their metabolic programs to ensure adequate supply of nutrients. Accumulating evidence suggests that changes in metabolism can also regulate shifts in cell fate [39,89]. Importantly, metabolism can do more than supply biosynthetic substrates - it actively modulates the signaling pathways that drive differentiation, positioning it as both a consequence and regulator of cell fate transitions [87,89]. Multiple regulators of cellular metabolism pathways that affect proliferation and differentiation were also differently expressed between the regenerative and non-regenerative regions in *C. robusta*. The PI3K/ AKT/mTOR pathway is a major pathway induced by growth factors, which promotes anabolic pathways and cell proliferation in multiple contexts [38,39,88,89]. Members of these pathways, including CASTOR2 [37], TSC2 [40], PTEN [41], PIK3R5 [39] as well as insulin growth factors (IGF1) [42] and transforming growth factor beta (TGF-β) [39], showed distinct expression patterns between the two body sections (Figs 5B, 5C and 5E). Interestingly, this pathway also controls stem and progenitor cell fate [89]. Loss of PTEN, a negative regulator of the pathway, leads to changes in stem cells proliferation [41,90], while deletion of TSC, a negative regulator of mTOR, depletes stem cells [40].

This divergence in metabolic related process was also in late regeneration stages (Figs 6B - 6C). At these stages, the regenerative body part showed an anabolic related process which includes differentiation and reorganization of the ECM, while the upper body part continues to show stress response that includes starvation and increased apoptosis (Fig 6B). The sustained differences in metabolic related pathways, which govern metabolism, proliferation, and cell fate decisions, between the two regions of *C. robusta* suggest that spatially distinct signaling environments play a decisive role in guiding cells to adopt regenerative or non-regenerative fate.

### Accumulating phagocytes at the wound site play distinct roles in regenerative and non-regenerative tissues

Additional variation in signaling environment between the two body regions can be seen in the immune-related factors found at each region. Following injury, phagocytes are accumulating at the wounded area in both body sections (Figs 7C and 7D). Yet, our transcriptomic data shows that the factors regulating the state and activation of these cells such as NLRP3 and NFE2L2 [63], vary between the regenerative and non-regenerative tissues (Fig 7I), thus affecting the overall regenerative outcome. It is known that inflammatory phagocytes are engaged in the initial response to wounding, and alternatively activated phagocytes are essential for wound closure and the resolution of tissue repair[56]. Their functional phenotype is dependent on the wound microenvironment, which changes during healing, hereby altering phagocytes phenotype [56,88,91]. Among the well-studied factors that involved in the temporal activation and transition of such phagocytes is the nuclear factor erythroid 2-related factor 2 (NRF2) [63,65,66]. The activation of this transcription factor can have antioxidant and anti-inflammatory effects through signaling transductions including JAK/STAT signaling pathways, with the regulation or cooperation of NLRP3 and MAPK signaling [63]. Both NFE2L2 (NRF2) and NLRP3 were upregulated in the regenerative body region in *C. robusta* (Fig 7I) suggesting a temporal transition in this cell response following injury.

### Conserved regeneration-associated genes exhibit region-specific expression in *Ciona robusta*

This study provides the first comprehensive transcriptomic roadmap of successful versus failed regeneration in *C. robusta*, highlighting JAK/STAT and MAPK signaling as key signatures of a pro-regenerative environment (Figs 6C - 6D and S5 Fig). As these correlations do not establish causality, future functional studies will be critical to determine their mechanistic roles in driving regeneration.

The JAK/STAT pathway is a conserved metazoan signaling system that transduces cues from extracellular cytokines into transcriptional changes in the nucleus [92]. It regulates important processes in stem cells, as well as regenerative processes in a conserved way between invertebrate and vertebrates [50,52,92–94]. Repressing this pathway leads to reduced proliferation and decreased survival of multiple cell lineages in mice [95,96]. In zebrafish, this pathway is crucial for the regeneration of diverse cells and tissue types, including cardiomyocytes, the retina and inner ear hair cells [97–99]. Interestingly, this pathway was also found to interact with MAPK signaling [94,100,101]. The mitogen-activated protein kinase (MAPK)/extracellular signal-regulated kinase (ERK) pathway have a central role in regenerative processes [102]. This pathway is activated following injury, orchestrating pro-regenerative responses such as promoting cell survival, regulating cell fate transitions, supporting migration and proliferation, and enhancing transcriptional and translational programs [102]. JAK1 was consistently upregulated in the regenerating body region of *C. robusta* across all time points (S5 Fig). Additional components of the JAK/STAT and MAPK signaling pathways also displayed distinct expression profiles between the two body parts, suggesting region-specific pathway activation (Figs 6C - 6D). Gene ontology analysis further revealed a temporal difference in ERK pathway repression, occurring earlier in the non-regenerating region compared to the regenerating counterpart (Fig 5C). Together, these findings point to a potential role for JAK/STAT and MAPK signaling in orchestrating the regenerative response in *C. robusta*.

### Distinct stem cell dynamics following injury underlie regenerative outcomes

The integration of single-cell and bulk transcriptomic analyses provides a comparative framework for understanding the cell type and molecular dynamics that facilitate injury response of the two body fragments. Our findings indicate that following injury, stem and progenitor cell populations exhibit distinct dynamic behaviors in regenerative versus non-regenerative contexts, reflecting differences in their wound response and their capacity to initiate and sustain the cellular trajectories required for regeneration.

The regenerative bottom fragment exhibits pronounced and coordinated temporal dynamics, with early changes in stem-associated populations followed by expansion of progenitor populations and subsequent enrichment of specialized cell types. These sequential transitions are consistent with activation of a regenerative program involving stem cell activation, proliferation, and progressive differentiation required for tissue reconstruction. In contrast, the non-regenerative upper fragment displays more limited and less coordinated changes in stem and progenitor populations over time. While some level of activation is detectable, the magnitude and progression of these responses are reduced, suggesting an incomplete or insufficient activation program. This impaired response occurs despite the presence of comparable stem cell populations, indicating that the limitation lies not in cell availability but in their activation state and ability to sustain a regenerative response.

Our study uses a comparative approach to define the cellular and molecular processes that facilitate regeneration in *Ciona robusta*, enabling the identification of key factors that distinguish regenerative from non-regenerative outcomes. By isolating an enriched putative stem cell population using ALDH as a non–species-specific marker and profiling it at single-cell resolution, we identified candidate stem cell markers and their spatial distribution following injury, enabling the tracking of stem cell dynamics across regenerative and non-regenerative fragments. Integration of these data reveals that regenerative outcomes are specified early after wounding, as tissue-level responses initially shared between fragments rapidly diverge into distinct molecular programs, despite the presence of similar putative stem cell populations. These programs include region-specific activation of immune, metabolic, proliferative, and differentiation pathways, with the regenerative fragment characterized by activation of key signaling pathways such as JAK/STAT and MAPK. Together, these findings indicate that the local signaling environment is a key determinant of regenerative success, governing whether cells engage in a productive regenerative trajectory. More broadly, this comparative framework highlights conserved principles of regeneration across chordates and suggests that successful regenerative strategies will require both stem cell competence and modulation of the surrounding niche to enable coordinated tissue reconstruction.

## Methods

### Animal husbandry

*C. robusta* adults were obtained from M-Rep (San Diego, California) and transferred to the mariculture facility at Hopkins Marine Station where they were maintained in seawater at 18°C. Gametes for fertilization were obtained based on previously published protocols[103]. Following fertilization, juveniles were cultured at the mariculture facility for future experiments.

### Transgenic line

Transgenic juvenile animals ubiquitously expressing the GFP reporter from a promoter of the gene Elongation Factor-1 alpha (EF1α::GFP) were obtained from the *C. robusta* Transgenic Line Resource center (CITRES, http://marinebio.nbrp.jp/ciona/).

### FACS analysis

Cell suspensions from whole transgenic animals were generated by mechanical dissociation using insulin syringe plunger and filtered through a 70-μm mesh followed by a 35-μm mesh into a 5 ml tube containing serum-free medium. No enzymatic dissociation was carried out. Cells were centrifuged at a speed of 500 G for 5 min, supernatant was discarded, and cells were resuspended in 1 ml of fresh medium. To enrich for stem cells, 1 μl of ALDH (AldeRed ALDH Detection Assay, SIGMA) was added. Cells were incubated for 30 min on ice under dark conditions. After gating for GFP positive cells, we analyzed the cells using forward (FSC) and back-scatter (BSC), and fluorescence from the ALDH staining using a Sony MA900 FACS instrument (Figs 1A and 2A). For morphological observations, the sorted cells were collected in 5 ml tubes containing 1 ml medium and centrifuged at a speed of 500 G for 5 min. The supernatant was discarded and the remaining medium (∼30 µl) was spun down, gently mixed and transferred to µ-slide 18 well glass bottom chamber (Ibidi). The chamber was placed on ice for 20 min to allow cells to settle down followed by imaging using Zeiss LSM700 confocal microscope (Fig 1A).

### In vivo transplantation experiments

For injection, glass needles were prepared using a P-1000 micropipette puller (Sutter Instruments). Needles were loaded with 5 μl of cell suspension (equivalent to ∼10,000 cells) and injected using manual micromanipulator (World Precision Instruments) under a stereoscope. All animals used for the transplantation experiment were derived from the same egg sample, collected from multiple adults, and mixed prior to fertilization with naïve or transgenic sperm sample. Cell suspension from a transgenic whole animal was generated using mechanical dissociation as described above. Cells with high ALDH activity from the FACS population (cStem sample) were sorted, centrifuged, resuspended in 500 µl of medium and injected under a stereoscope into the heart of recipient animals (n=26, 3 months old) (Fig 2A). The control recipient grope (n=17) received the same treatment with cells with low ALDH activity from the FACS population (Control sample). 24 hr post injection the oral siphon of all recipient animals was amputated using a scalpel. The animals were kept at running sea water and were monitored for survival and regeneration. 30 days post-transplantation, GFP+ cells were sorted from the animals in both treatments groups and captured for scRNA-Seq to test whether the transplanted cells survived and differentiated within the recipient body (Fig 2A).

### Single cell RNA sampling, library preparation, sequencing and alignment

Single cell capture and library preparation with 10x Chromium Next GEM Single Cell 3ʹ Reagent Kits v3.1 (10x Genomics) and the Chromium Controller (10x Genomics) were performed using the manufacturer’s protocol with slight modifications. 10,000 cells were sorted using Sony MA900 FACS into tube containing 2 μl of media and washed once in Calcium Magnesium Free Seawater (495mM NaCl, 9.7mM KCl, 27.6mM, NaHCO_3_, 50mM Tris-HCl pH8, filtered at 0.2µm), spun down at 4°C for 5 min at 500G and added to 10x Master Mix supplemented with 0.7D-Mannitol, for a final osmolarity of 770mOsm/L in each capture channel. cDNA amplification was performed with 11 cycles for ALDH, cStem, control and cStem 30dpt. For sample index PCR, 15 cycles were performed. Final libraries were quantified using a Qubit fluorometer and fragment sizes were assessed using a TapeStation with High Sensitivity D5000 tapes. Sequencing was performed on an Illumina NovaSeq 6000 platform with paired-end 150 bp reads, achieving a depth of approximately 160–330 million reads per library, using 28 cycles for Read 1, 10 cycles for each i5 and i7 indices, and 90 cycles for Read 2. Raw sequencing reads were demultiplexed and aligned to the Kyoto 2021 gene model (KY21)[104] using the CellRanger count pipeline 7.1.0, following standard parameters.

### Quality Control and Data Integration

Initial quality control was performed to exclude low-quality cells and potential doublets. The raw dataset contained 18827 features across 11351 cells. Cells were annotated with their sample of origin based on cell barcodes. The percentage of mitochondrial transcripts per cell was calculated using the Seurat function ‘PercentageFeatureSet’ with the pattern “Mitochondria”. Cells were filtered based on the number of detected transcripts, the number of detected genes, and the percentage of mitochondrial transcripts. The lower bounds are set to zero and upper bounds defined as the median plus 5 times the median absolute deviation (MAD) for both metrics. Genes expressed in more than five cells were retained for further analysis. After filtering, the final dataset contained 14144 features across 8465 cells. Data integration was performed using Seurat’s integration workflow, aligning cells across samples to correct for batch effects. The integrated dataset was normalized using the SCTransform workflow, regressing out percent mitochondrial content, number of features, and number of counts per cell to minimize technical variation.

### Cell Cycle Analysis

Cell cycle scoring was performed using a custom set of S phase and G2/M phase marker genes. Specifically, the following genes were used for S phase: PCNA, MCM2, RRM1, UHRF1, RPA2, NASP, SLBP, POLD3, RRM2, CDC45, CDC6, TIPIN, CASP8AP2, POLA1, and CHAF1B. For G2/M phase, the markers included NUSAP1, TPX2, CKS2, MKI67, TACC3, SMC4, KIF11, KIF20B, NCAPD2, DLGAP5, CDCA8, ANLN, LBR, CKAP5, CENPE, CENPF, TUBB4B.3, and ECT2. Cell cycle phase scores were calculated for each cell using Seurat’s ‘CellCycleScoring’ function, with the custom S and G2/M gene lists provided as input. Each cell was assigned an S score, a G2/M score, and a predicted cell cycle phase (G1, S, or G2/M).

### Dimensionality Reduction and Clustering

Principal component analysis (PCA) was performed on the integrated assay, computing 50 principal components. The optimal number of principal components for downstream analysis was estimated using the ‘intrinsicDimension::maxLikGlobalDimEst’ function, with a cutoff of 10 PCs. Clustering was performed by constructing a nearest neighbor graph using the first 12 principal components, followed by community detection with a resolution parameter of 0.4. The quality of clustering was evaluated using silhouette scores 0.43. Dimensionality reduction for visualization was achieved using UMAP, projecting cells into a two-dimensional space based on the first 10 principal components.

### RNA Velocity and Pseudotime Analysis

RNA velocity analysis was performed to infer cellular dynamics and developmental trajectories using scVelo and velocyto. Spliced and unspliced transcript counts were generated from the aligned BAM files using velocyto run10x, producing spliced and unspliced count matrices. Velocity analysis was performed using scVelo’s dynamical model workflow. The merged dataset was filtered and normalized using ‘scv.pp.filter_and_normalizè with a minimum of 30 shared counts and selection of the top 3,000 highly variable genes. Moments were computed using 30 principal components and 30 nearest neighbors. RNA velocity was estimated using the dynamical model (’scv.tl.recover_dynamics’ and ‘scv.tl.velocity’ with mode=’dynamical’), which models the full transcriptional dynamics including gene-specific reaction rates Velocity graphs were constructed to represent the cell-to-cell transition probabilities, and latent time was recovered using ‘scv.tl.recover_latent_timè to provide a measure of pseudotime based on the underlying gene expression dynamics.

### Cytotrace2 Analysis

To assess cellular differentiation potential, Cytotrace2[105] was applied to the filtered and integrated dataset. This analysis provided a quantitative measure of differentiation state for each cell, facilitating the identification of progenitor and differentiated populations.

### Bulk RNA-seq Deconvolution Analysis

Cell type composition was estimated from bulk RNA-seq data using scRNA-seq reference-based deconvolution. A total of 38 bulk samples across 10 unique conditions (upper and bottom body fragments at 0, 4, 12, 24, and 72 hours post-bisection) were deconvolved into cell stages (stem, progenitors, specialized as defined in Fig 1G) and 20 cell types using Bisque [106]. Method performance was assessed by comparing deconvolved proportions to reference scRNA-seq cell composition. Pearson correlations of 1.0 for cell stages and 0.832 for cell types. Cell types with absolute error >0.05 or relative error >0.5 were excluded, resulting in removal of cell type clusters 7 and 19.

### Software and Code Availability

All analyses were performed in R (4.4.2) and Python (3.6.1) running on CentOS Linux 7. Key packages used included Seurat (5.3.0), SingleCellExperiment (1.28.1), CytoTRACE2 (1.1.0), scanpy (1.7.2), scvelo (0.2.5), velocyto (0.17.16).

### Regeneration Experiment

*C. robusta* juveniles of the same age (3 months, 0.5-1 cm in length) were anesthetized (Tricaine, Sigma) and bisected using a scalpel (Fig 3A). Both treated and non-amputated animals were maintained in running seawater for the duration of the experiment. Response to touch was used to determine survival.

### Bulk sequencing

Upper and bottom body sections of *C. robusta* (n = 24, 3 months old) were collected at six different time points following injury-immediately following amputation (0 hpa), 4, 8, 12, 24 and 72 hours post amputation (n=4 per time point). Total RNA was prepared using Trizol (Life Technologies) following the manufacturer’s directions. Tissues were homogenized in Trizol (1 ml) using TissueLyser II and Tungsten Carbide Beads, 3 mm (Qiagen). Following homogenization, the samples were chloroform-extracted, precipitated with isopropanol, washed with ethanol and dissolved in sterile, RNase-free water. Samples were analyzed on an Agilent 4150 TapeStation to determine quality prior to library preparation. cDNA and libraries were prepared using NEBNext Single Cell/Low Input RNA Library Prep Kit and barcoded using NEBNext Multiplex Oligos for Illumina. All magnetic bead purification was accomplished using DNA Spri Beads. Samples were then analyzed on an Agilent 4150 TapeStation to determine the concentration of each sample prior to sequencing. On average, 20 million 2x150 bp reads (NovaSeq X Plus) were sequenced for each library.

### Bulk sequencing analysis

Read mapping was performed as previously described[107]. Briefly, sequences were demultiplexed followed by quality trimming with TrimGalore v0.5.0 and aligned to the *C. robusta* genome [“Ghost” Kyoto Ciona genome[108]] with STAR program[109] (parameters: --outFilterMultimapNmax 20 --outFilterMismatchNoverLmax 0.04 --alignSJoverhangMin 8 --alignSJDBoverhangMin 1 --sjdbScore 1 --alignIntronMax 1,000,000 --alignMatesGapMax 1,000,000). The bam files from the STAR alignment were used to calculate expression levels of genes and transcripts and incorporate them into a gene count table using HTSeq v2.0.1_py39 [110].

The gene count table was then used to perform differential gene expression analysis using DESeq2 [111]. Heatmap plots were generated to visualize gene expressions across samples using the ‘pheatmap’ package in R. Before generating the heatmaps, the raw gene counts were normalized using the log2(CPM) method. A small constant (0.1) was added to all the values before log transformation since log(0) is undefined. Therefore, the lowest value in the normalized count table is log2(0.1) = -3.32.

Sets of genes were tested for enrichment of Gene Ontology (GO Biological Process, Molecular Function and Cellular Component) terms. For a set of genes with significant up and down effects, found using the Wald test, an over-representation analysis (ORA) was performed using the enricher function of the clusterProfiler package (v3.16.1) in R with default parameters [112]. A gene set enrichment analysis (GSEA) was also performed on the entire assembly using a scoring based on the log fold change of each gene on its respective time point using the GSEA function of clusterProfiler package (v3.16.1) in R with default parameters [112]. In both cases Significant GO terms were identified with an FDR < 0.05. The gene ontology terms were obtained from a manually created database based on the SwissProt curated *Mus musculus* GO annotations, using the makeOrgPackage function of AnnotationForge [113].

### In situ hybridization

Short in situ hybridization antisense DNA probes were ordered as lyophilized oligo pools from Integrated DNA Technologies and were resuspended in nuclease-free water to a concentration of 0.5 µM. For each gene, the probe was designed based on the split-probe design of HCR v.3.0 [114] using HCR 3.0 Probe Maker v0.3.2 [115] with adjacent B1 amplification sequences (S11 Table).

For Hybridization Chain Reaction (HCR) in situ hybridization, *C. robusta* samples (n=12, 3 months old) were incubated in fixation buffer (4% PFA in 1x PBS) overnight at 4 °C, followed by 100% methanol dehydration and storage at −20 °C for at least 24 h. The samples were gradually rehydrated in 75%, 50% and 25% methanol diluted with PBS (5 min for each step) followed by two washes in 1x PBS for 5 and 10 min, respectively. The samples were permeabilized for 30 min at room temperature in detergent solution (1% SDS, 0.5% Tween-20, 50 mM Tris-HCl (pH 7.5), 1 mM EDTA (pH 8), 150 mM NaCl). The samples were then washed 2 × 5 min with PTw (0.1% Tween-20 in 1x PBS) followed by post-fixation in 4% PFA for 25 min and washed again 3 × 5 min with PTw. The samples were prehybridized in hybridization buffer (Molecular Instruments) for 30 min at 37 °C. The probes were then added to the hybridization buffer at a final concentration of 0.02 µM and the samples were allowed to hybridize at 37 °C for overnight but no more than 16 h. Following hybridization, the samples were washed 2 × 30 min in probe wash buffer (Molecular instruments) at 37 °C and then for 5 min in 5x SSCT (5x sodium chloride sodium citrate (SSC), 0.1% Tween-20) at room temperature. They were then pre-amplified in amplification buffer (Molecular Instruments) for 30 min. At the same time, H1 and H2 components of the HCR hairpins B1 coupled to either 546 or 647 amplifiers (Molecular Instruments) were incubated separately at 95 °C for 90 s, cooled down to room temperature in the dark for 30 min and then pooled together before being added to the amplification buffer at a final concentration of 30 nM. The amplification was then performed at room temperature for overnight but no more than 16 h. The samples were subsequently washed 2 × 5 min and additional 2 × 30 min in 5x SSCT followed by incubation for 2 h in PBS containing 0.8:500 DAPI. Finally, the samples were washed 3 × 10 min in PBS, transferred to 50% glycerol for at least 1 h and then mounted on a glass slide for imaging. Imaging was done using Zeiss LSM700 confocal microscope. For each sample, a series of optical sections were taken with a *z*-step interval of 3–5 µm. Multichannel acquisitions were obtained by sequential imaging and the confocal optical sections were processed using ImageJ v.1.54j [116].

### Antibodies staining

Juveniles (n = 12, 3 months old) were bisected and fixed at three time points (Amputation, 24 and 72 hours post amputation, Fig 3B) using 4% paraformaldehyde in phosphate-buffered saline (PBS) overnight at 4°C. Fixed upper and bottom samples were then washed three times (5 min each) in PBS followed by a 10 min wash in 0.5% TritonX-100 at room temperature (RT). Samples were blocked in PBT + 3% BSA for 3 h at RT. Antibodies were added directly to the blocking solution overnight at 4°C. To visualize nerves and cilia, we used a mouse monoclonal anti-acetylated tubulin antibody (Sigma T7451) diluted 1:1000 in blocking solution (36, 3, 7). Samples were then washed twice for 10 min each in PBT. Secondary antibodies (Alexa Fluor goat anti-mouse IgG 488, ThermoFisher, Waltham, Massachusetts, A-11001) were added at 1:500 dilution to the blocking solution for 3 h at RT. Samples were then washed in PBS three times for 10 min each. Samples were stained with DAPI (Sigma-Aldrich, Missouri, 1 μg/mL in PBSTx) and mounted. Images were taken using Zeiss LSM700 confocal microscope.

### EdU staining

Cell proliferation was detected by incorporating 5-ethynyl-2-deoxyuridine (EdU) into replicating DNA. *C. robusta* juveniles (n=9, 3 months old) were divided into three groups, bisected and separated into upper and bottom body sections using a scalpel (Figs 4D - 4E). The first group (*n* = 4) was exposed to a 4 h EdU pulse following amputation and was fixed immediately afterwards. The second group (*n* = 4) was exposed to a 4 h EdU pulse 20 h following amputation and was fixed immediately afterwards and the third group (*n* = 4) was exposed to a 4 h EdU pulse 68 h following amputation and was fixed immediately afterwards (Fig 4E).

For the pulse experiments animals were incubated with 10 μmol/L EdU (Invitrogen, Carlsbad, California) in 5 mL of MFSW for 4 h in Petri dishes. Following completion of the labeling, animals were fixed for 12 h in 4% FA, rinsed three times in 1 × phosphate-buffered saline (PBS), and processed for EdU detection using Alexa Fluor azide 488 at room temperature, according to the instructions of the Click-iT EdU Alexa Fluor High Throughput Imaging Assay Kit (Invitrogen). Samples were stained with DAPI (ThermoFisher 33342) (1 μg/mL in PBS) and mounted in VECTASHIELD (Vector Laboratories RK-93952-28) using coverslips.

### Phagocytosis assay

pHrodo Deep Red E. coli BioParticles Conjugate for Phagocytosis were reconstituted in 100 μl of phosphate buffered saline (PBS), according to the manufacturer’s specifications (Invitrogen) and stored at 4 °C until needed. For injection, glass needles were prepared using a P-1000 micropipette puller (Sutter Instruments). Needles were loaded with 5 μl of the bioparticle stock and injected using manual micromanipulator (World Precision Instruments) under a stereoscope into the heart of recipient animals (n=8, 3 months old). Successful microinjection was determined by visualizing extension of the heart. Imaging was done using Zeiss LSM700 confocal microscope. For each sample, a series of optical sections were taken with a *z*-step interval of 3–5 µm. Multichannel acquisitions were obtained by sequential imaging and the confocal optical sections were processed using ImageJ v.1.54j [116].

### Statistical Information

Detailed description of the statistical method and number of samples used in the different experiments is included in the main text, figures, and methods. No statistical methods were used to predetermine sample size.

## Supporting information

supplemental images

## Data availability

The sequencing data generated during this study is available in NCBI. The scRNA-Seq data is under accessions: PRJNA1313632. The code used for the scRNA-Seq analysis are available on GitHub at https://github.com/chstellar/Regional-Signaling-Controls-Stem-Cell-Mediated-Regeneration-in-an-Invertebrate-Chordate. The bulk RNA data is under accessions: PRJNA1328382.

## Acknowledgement

We thank Dr. Omri Wurtzel for feedback and discussions of the work, Prof. Chris Lowe and Prof. Larry Crowder for sharing laboratory equipment and lab space, Katherine J. Ishizuka and Karla J. Palmeri for assistance with animal culture, Prof. Yasunori Sasakura for his support and assistance with obtaining transgenic animals, Prof. Emma Farley for assistance with fertilization protocols.

## Supporting information

**Fig S1. Gene markers used for cluster annotation.** (A) Dot plot showing representative gene markers per cell cluster based on scRNA-seq analysis of *Ciona robusta*. Leiden cluster numbers are indicated. Data is correlated with Supplementary Tables 1 and 2. (B) UMAP expression plots of selected markers for stem cells, neural cells, epithelial cells, and immune cells based on scRNA-seq data. Data is correlated with S1 and S2 Tables.

**S2 Fig. Gene markers per cell cluster.** Heatmap showing the top 5 gene markers per cell cluster according to scRNA-Seq of *C. robusta*. Leiden numbers are indicated. Data is correlated with S2 Table.

**S3 Fig. Morphological differentiation of candidate stem cells after transplantation.** Flow cytometry analysis of forward scatter (FSC, cell size) and side scatter (SSC, granularity/complexity) on a logarithmic scale. Shown from left to right: cStem population (GFP+) before transplantation from EF1α::GFP, cStem (GFP+) 30 days post-transplantation in WT animals, and total control GFP+ population in EF1α::GFP animals.

**S4 Fig. Injury induces systemic proliferation and local accumulation of proliferating cells at the wound site in both regenerative and non-regenerative tissues.** (A–C) Whole-mount immunofluorescence showing higher-magnification views of the injury site in both body parts following bisection at (A) 4 hpa, (B) 24 hpa, and (C) 72 hpa. For each time point, the left panels show merged images of DAPI (gray) and EdU (green), while the right panels show EdU staining alone to highlight proliferating cells. (D-J) Quantification of EdU-positive cells in 100 µm2 sections (n = 3 animals per time point, 4 sections per animal). Box plots display the number of EdU-positive cells in each section. Data are mean. ^∗^p < 0.05; ^∗∗^p < 0.01; ^∗∗∗^p < 0.001. (D-E) Number of EdU-positive cells in tissue proximally and distal from the injury site in the anterior body parts (D) and posterior body part. (E). Two-tailed, t test. (F) Number of EdU-positive cells in the area distal to the injury site in both body parts at different time points following injury. Two-tailed, t test. (G-H) Number of EdU-positive cells in the anterior body part at different time points following injury. (G) Proximal to the injury site. (H) Distal from the injury site. Kruskal-Wallis test. (I-J) Number of EdU-positive cells in the posterior body part at different time points following injury. (I) Proximal to the injury site. (J) Distal from the injury site. Kruskal-Wallis test.

**S5 Fig. Comparative gene expression profiles of regenerative and non-regenerative tissues.** Expression profiling of significantly differentially expressed genes across all injury time points, comparing the regenerative bottom body part to the non-regenerative upper body part. Rows represent genes, columns show summary of gene expression in each time point. Color scale shows the log fold change, blue and red, low and high expression, respectively. Colored square on the left side of the heatmap represents gene function based on GO annotation.

**S1 Table. Gene markers used for cluster annotations showing function and reference.** Data is correlated with Figs 1, 2, and S1 Fig.

**S2 Table. Rank of gene expression for each leiden cluster from *C. robust* scRNA-Seq**. Data is correlated with Figs 1, 2, 3, and S2 Fig.

**S3 Table. GFP+ cells count data.** Data is correlated with Fig 2.

**S4 Table. Rank of differentially expressed genes from the upper body part at different time points following amputation.** Data is correlated with Figs 4, 5, 6, and 7.

**S5 Table. Rank of differentially expressed genes from the bottom body part at different time points following amputation.** Data is correlated with Figs 4, 5, 6, and 7.

**S6 Table. EdU count data.** Data is correlated with Figs 4 and S3 Fig.

**S7 Table. Rank of differentially expressed genes between both body parts at different time points following amputation.** Data is correlated with Figs 4, 5, 6, and S4 Fig.

**S8 Table. GO annotations for the upper body part at different time points following amputation.** Data is correlated with Figs 5 and 6.

**S9 Table. GO annotations for the bottom body part at different time points following amputation.** Data is correlated with Figs 5 and 6.

**S10 Table. GO annotations for differentially expressed genes between both body parts at different time points following amputation.** Data is correlated with Fig 5.

**S11 Table. Short in situ hybridization antisense DNA probes used in this study.** Probes were designed with adjacent B1 amplification sequences.

**S12 Table. PhRodo count data.** Data is correlated with Fig 7.

