## Supplementary figures and images for "Regional Signaling Controls Stem Cell-Mediated Regeneration in an Invertebrate Chordate"

### Figure_S1.jpg

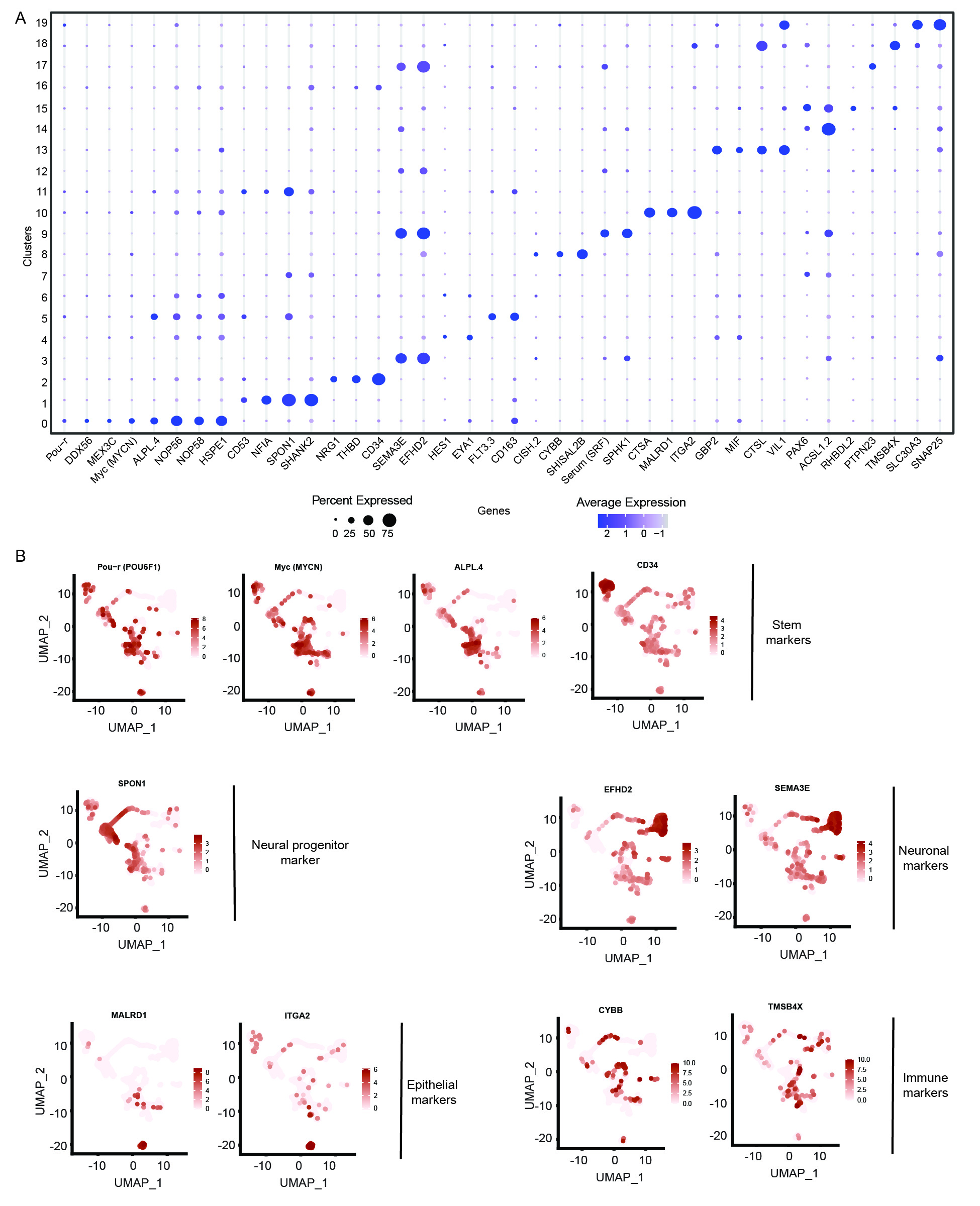

### Figure_S2.jpg

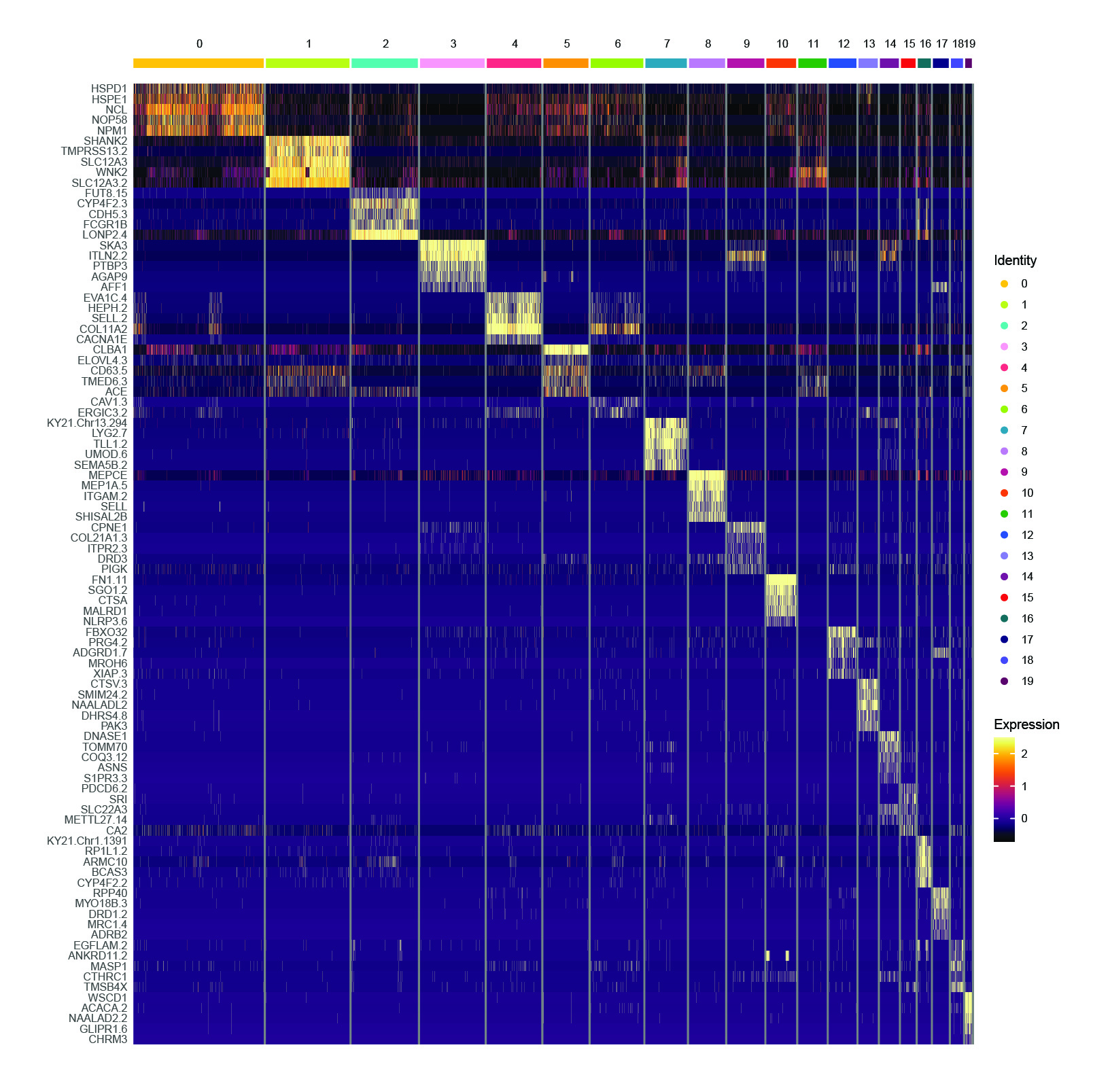

### Figure_S3.jpg

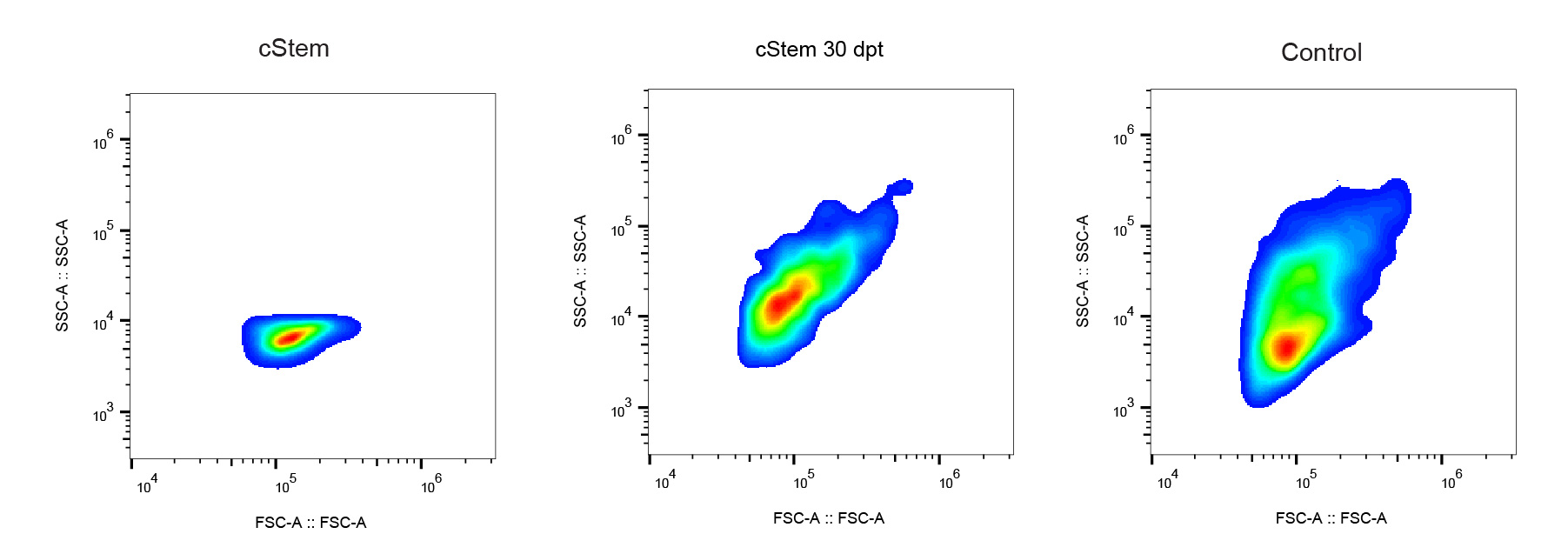

### Figure_S4.jpg

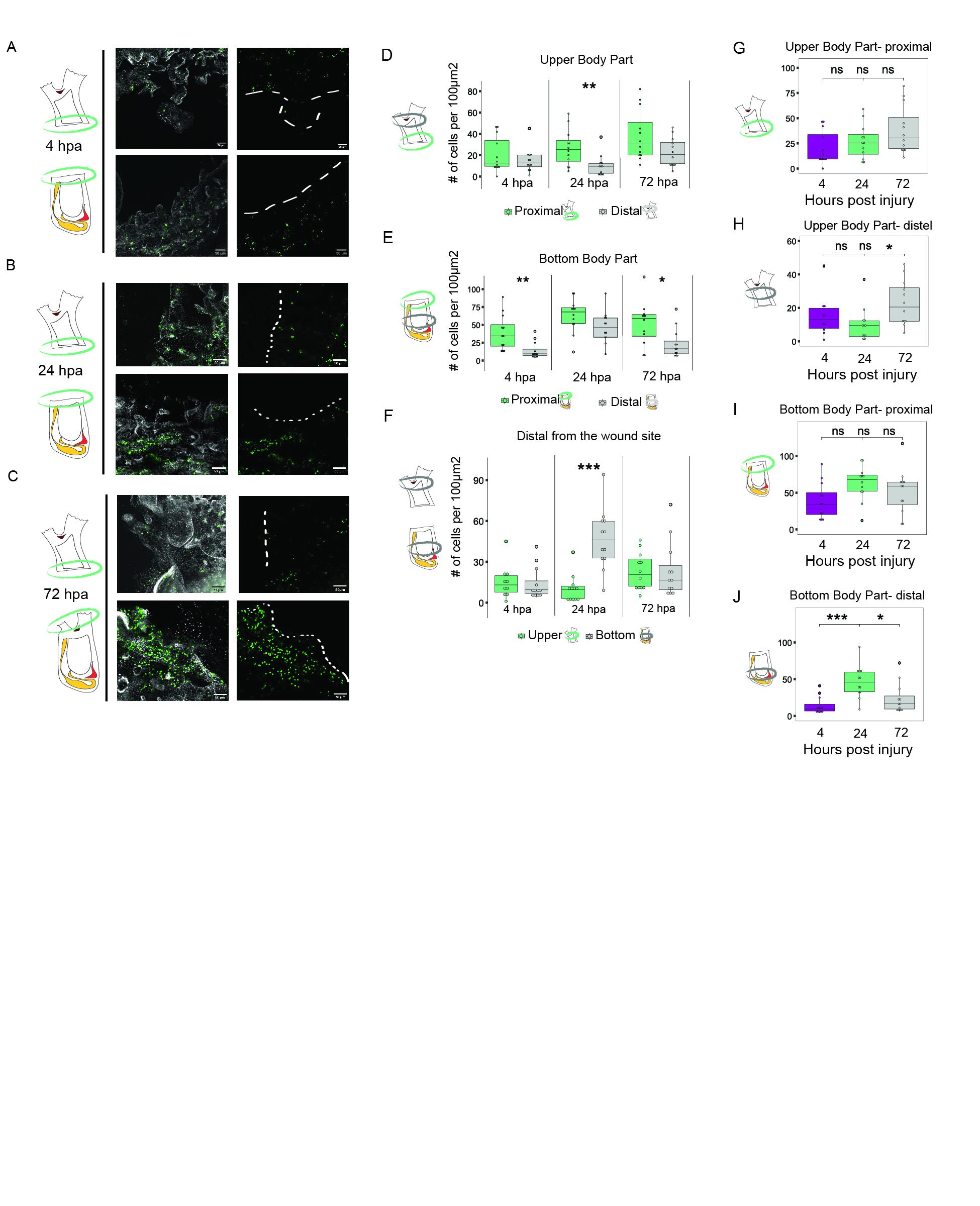

### Figure_S5.tif

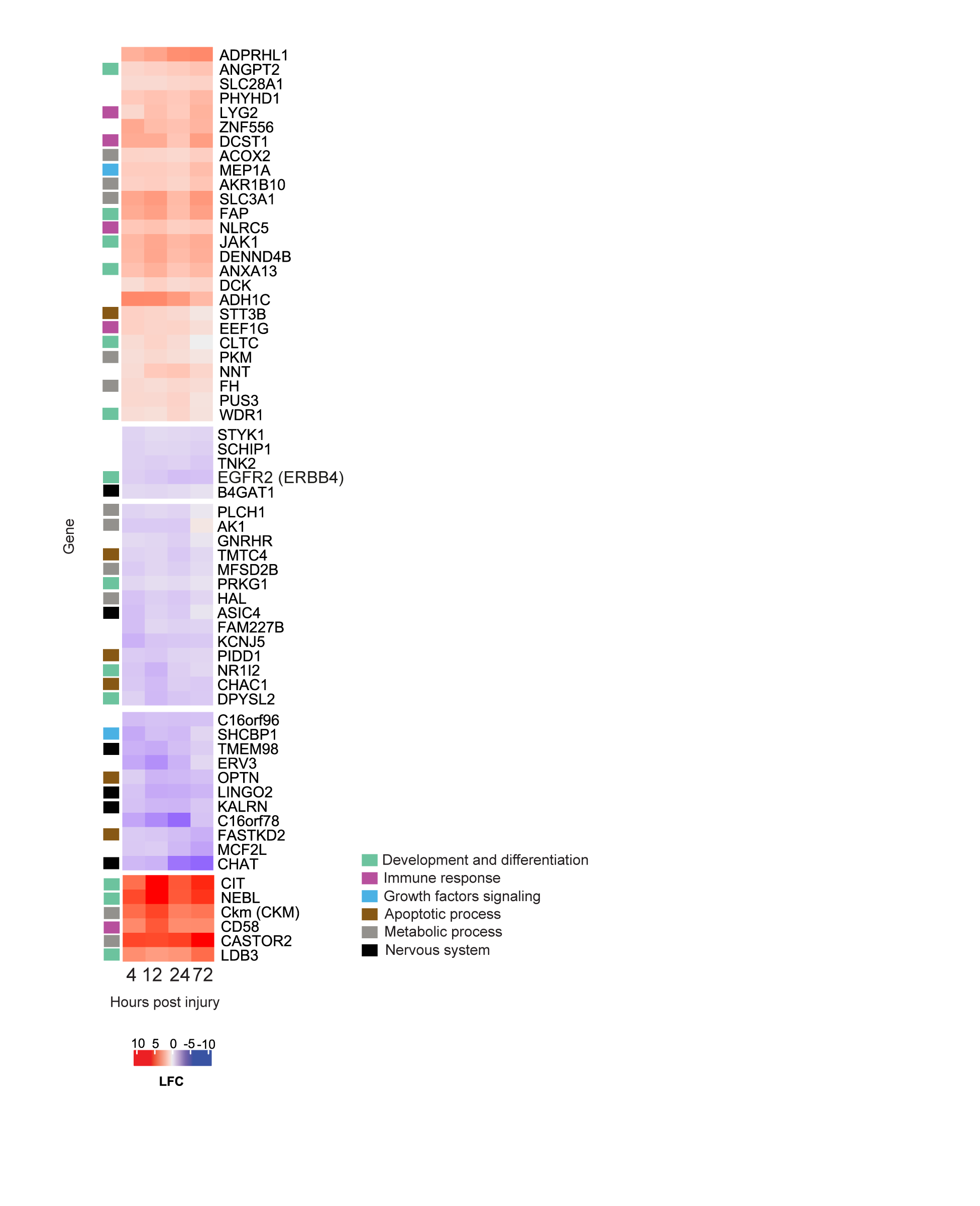
